# Excess salicylic acid mediates graft incompatibility by repressing auxin-dependent vascular reconnection in tomato

**DOI:** 10.64898/2026.07.30.741952

**Authors:** Ruiduo Han, Kaiwei Meng, Xiujuan Wang, Kaili Mao, Zefeng Chen, Xiaojian Xia, Huijia Kang, Yanhong Zhou, Hannah Rae Thomas

**Affiliations:** Department of Horticulture, Zijingang Campus, Zhejiang University, 866 Yuhangtang Road, Hangzhou 310058, PR China; Key Laboratory of Horticultural Plants Growth and Development, Chinese Ministry of Agriculture and Rural Affairs, Yuhangtang Road 866, Hangzhou 310058, PR China; Hainan Institute, Zhejiang University, Sanya 572025, PR China; Zhejiang Provincial Key Laboratory of Horticultural Crop Quality Improvement, Zhejiang University, Zijingang Campus, Hangzhou 310058, China

**Author notes:** Corresponding author, Hannah Rae Thomas.

**Keywords:** plant grafting, graft compatibility, salicylic acid, auxin, tomato

## Abstract

Graft incompatibility limits the combination of scions and rootstocks, yet the signals that block vascular reconnection remain poorly understood. Using incompatible tomato-pepper (*Solanum lycopersicum-Capsicum annuum*) grafts, we found that salicylic acid (SA) overaccumulates at the graft junction, predominantly in the rootstock, while incompatible scions exhibit enhanced pattern-triggered immunity. SA-deficient *nahG-*expressing tomato failed to rescue incompatibility and instead showed severely impaired self-graft healing, indicating that successful tissue reunion requires an optimal, rather than minimal, SA response. Defense-associated genes remained activated in failed grafts regardless of SA accumulation, placing SA downstream of incompatibility determination. Exogenous SA application phenocopied incompatibility, blocking xylem reconnection, inducing cell death, and suppressing auxin signaling. Incompatible grafts similarly displayed reduced auxin accumulation and response at the graft junction. Exogenous auxin partially rescued xylem reconnection in incompatible grafts but failed to activate cambial regulator genes or fully restore compatibility. We propose that graft compatibility depends on SA-auxin homeostasis, where moderate SA supports wound-associated defense and regeneration, whereas excessive SA suppresses the auxin-dependent program required for xylem differentiation. These findings identify SA as a dose-dependent regulator of graft healing in tomato and link excessive immune activation to failed vascular regeneration.

## Introduction

Plant grafting is a horticultural technique that combines shoots, known as scions, with root systems, called root(stocks). While traditionally used on woody plants, it is now commonly applied to propagate vegetables, such as Solanaceous species (Mudge *et al*., 2009; Lee *et al*., 2010). Through grafting, stress-resistant rootstocks can be paired with valuable commercial scions to introduce advantageous traits without the need for genetic modification or slow traditional breeding. For example, wild tomato (*Solanum habrochaites*) rootstocks are often grafted with domesticated tomato (*Solanum lycopersicum L.*) scions due to their resistance to stress and disease (Mattos *et al*., 2011; Vanlay *et al*., 2022).

However, not all plants can be grafted together, a limitation known as graft incompatibility (Andrews *et al*., 1993). For a graft to be compatible, both the nonvascular and vascular tissues must reconnect (Thomas *et al*., 2023). Even closely related species can show graft incompatibility. (Zeist *et al*., 2017; Lee *et al*., 2024). A prime example of this is in tomato-pepper incompatible grafts, where, despite nonvascular tissue forming at the graft site, xylem files fail to reconnect (Thomas *et al*., 2022).

Xylem reconnection is controlled by a complex network of vascular regulators. A key component of this process is auxin, which is transported down the scion to the graft site through polar auxin transport (PAT) (Melnyk *et al*., 2015). The location of auxin is the driving force behind the induction of xylem differentiation, both during development, wound healing, and grafting (Scarpella *et al*., 2006; Asahina *et al*., 2011a; Fàbregas *et al*., 2015; Matsuoka *et al*., 2016). This is especially true during grafting, where treatment with auxin transport inhibitors can completely block graft healing, but can be rescued by local auxin treatment (Matsuoka *et al*., 2016).

Similarly, auxin signaling mutants, *ABERRANT LATERAL ROOT FORMATION 4 (*ALF4) and *BODENLOS* (BDL), have significantly reduced graft healing rates in Arabidopsis, highlighting the integral role of auxin in vascular reconnection (Melnyk *et al*., 2015; Serivichyaswat *et al*., 2024). Despite a clear physiological breakdown in xylem formation between the scion and stock of tomato-pepper grafts, the signals that inhibit the necessary intercellular communication required for xylem reconnection remain unknown.

However, recent work investigating the transcriptional profile of tomato-pepper genetic incompatibility found that tomato-pepper grafts also exhibit an upregulated and prolonged transcriptional signature enriched for defensive processes, including salicylic acid (SA) transcriptional responses (Thomas *et al*., 2024). This suggests that the basis for tomato-pepper genetic graft incompatibility may be due to an overactive immune response, but the role of SA in tomato graft incompatibility has not been investigated.

Plant grafting is a complex process partially regulated by hormones. While considerable attention has been paid to jasmonic acid (JA), a key regulator of wound response, salicylic acid has not yet been implicated in graft healing (Asahina *et al*., 2011b; Pitaksaringkarn *et al*., 2014; Ikeuchi *et al*., 2020; Matsuoka *et al*., 2021). While shown to accumulate during some wound responses (Sano *et al*., 1994; Liu *et al*., 2008; Ogawa *et al*., 2010; Ikeuchi *et al*., 2020), SA is primarily functional in response to biotrophic bacteria (Mishra *et al*., 2024). SA is a well-studied component of the defense response. Both pattern-triggered immunity (PTI) and effector-triggered immunity (ETI) elicit SA biosynthesis (Ding *et al*., 2018; Zhang & Li, 2019). Unlike many hormones, which possess a single receptor, SA can bind to NONEXPRESSOR OF PATHOGENESIS-RELATED 1 (NPR1), NPR3, NPR4, as well as numerous other proteins known as SA Binding Proteins (Uquillas *et al*., 2004; Fu *et al*., 2012; Pokotylo *et al*., 2019). In the absence of SA, NPR3/4 act as negative repressors of defense-related genes. SA binds and inhibits the transcriptional repression of NPR3/4. In parallel, SA binds to NPR1 to promote transcription of various defense genes, such as *Pathogenesis-related* (*PR*) proteins (Li *et al*., 2004; Powers *et al*., 2024). NPR1 also promotes additional SA biosynthesis via the ICS1 and PAL pathways (Fragnire *et al*., 2011). This network triggers the actions required for various immune processes, such as hypersensitive response (HR).

This complex and elegant mechanism is integral for maintaining the defense-growth balance. It has been previously shown that eliciting a strong immune response to pathogens, herbivores, or wounding comes at the cost of growth (He *et al*., 2022). Exogenous SA treatment or mutants with high SA levels exhibit dwarf phenotypes (Bowling *et al*., 1997; Rate *et al*., 1999a; Miura *et al*., 2010), while mutants with low levels of endogenous SA often have increased growth and yield (Scott *et al*., 2004; Abreu & Munné-Bosch, 2009). The mechanism that facilitates SA-mediated growth repression has been largely associated with hormone cross-talk. For example, SA represses growth via gibberellins by stabilizing DELLA proteins against degradation, thereby constraining cellular division (Li *et al*., 2019b). However, a larger body of evidence indicates that SA predominantly represses growth by inhibiting various auxin-related processes (Rawat & Laxmi, 2025). Since numerous pathogens possess mechanisms to stimulate plant growth via auxin, it is hypothesized that plants evolved a way to block auxin responses when SA was high (Glickmann *et al*., 1998). High SA can repress auxin biosynthesis, perception, transport, or signaling, depending on the species and concentration (Michniewicz et al., 2007; Wang et al., 2007a; Du et al., 2013; Armengot et al., 2016; Pasternak et al., 2019b; Kong et al., 2020; Han et al., 2025). One key way SA blocks overall plant growth is by inhibiting cell division and expansion (Vanacker et al., 2001; Miura et al., 2010). However, at low levels, SA positively regulates these processes (Rate *et al*., 1999b; Vanacker *et al*., 2001; Wang *et al*., 2021). This is supported by numerous studies that have shown that low-concentration exogenous SA application can stimulate, while high concentrations of SA block growth, suggesting that the role of SA moves between inhibiting and promoting growth in a concentration-dependent manner (Hayat *et al*., 2006; Kováčik *et al*., 2008; Canakci, 2011; Pasternak *et al*., 2019; Li *et al*., 2022).

Auxin-SA homeostasis is known to play a critical conserved role in balancing plant growth and defense priorities, yet the functional interplay of these two antagonistic hormones during graft compatibility and healing has remained almost entirely uncharacterized. In this study, we systematically investigated SA dynamics across both compatible tomato/tomato self-grafts and incompatible tomato-pepper heterografts, revealing a previously unreported dose-dependent role for SA in regulating graft healing outcomes. We demonstrate that moderate SA levels are required for normal graft healing, excessive SA accumulation disrupts auxin-mediated vascular reconnection, and the absence of SA inhibits tissue reconnection even in genetically compatible combinations.

## Results

### Incompatible rootstocks overaccumulate salicylic acid after grafting

Previously, the role of salicylic acid (SA) in graft healing and compatibility was poorly studied. To understand the involvement of this defense hormone, we used the tomato-pepper model system, which exhibits delayed graft incompatibility (Thomas *et al*., 2022, 2024). We first explored SA accumulation in compatible self-grafted tomato and self-grafted pepper. Previous work has shown that self-grafted tomato requires 5 days to form xylem reconnections, while pepper requires 7 days (Thomas *et al*., 2022). Following a similar trend, SA content steadily increased in both the scion and stock in compatible grafts until approximately 1 day prior to xylem reconnection, with total SA accumulation peaking in the stocks 4 days after grafting (DAG) in tomato and 6 DAG in pepper (Figures 1A-B). Both species exhibited peak SA accumulation in the scion 1 day prior to the peak accumulation in the stock, suggesting a temporal component to SA signaling. In both species, the stock showed greater accumulation than the scion, with 2.7 (pepper) and 2 (tomato) times higher SA, with the most significant differences occurring between the two tissues 7 DAG. This is an interesting finding, as SA is not classically associated with wound response, yet it has been shown to accumulate in certain scenarios (Liu *et al*., 2008). However, this is the first time that spatial accumulation of SA below a stem wound has been demonstrated, suggesting that SA may exhibit spatially regulated accumulation during graft healing.

**Figure 1.**
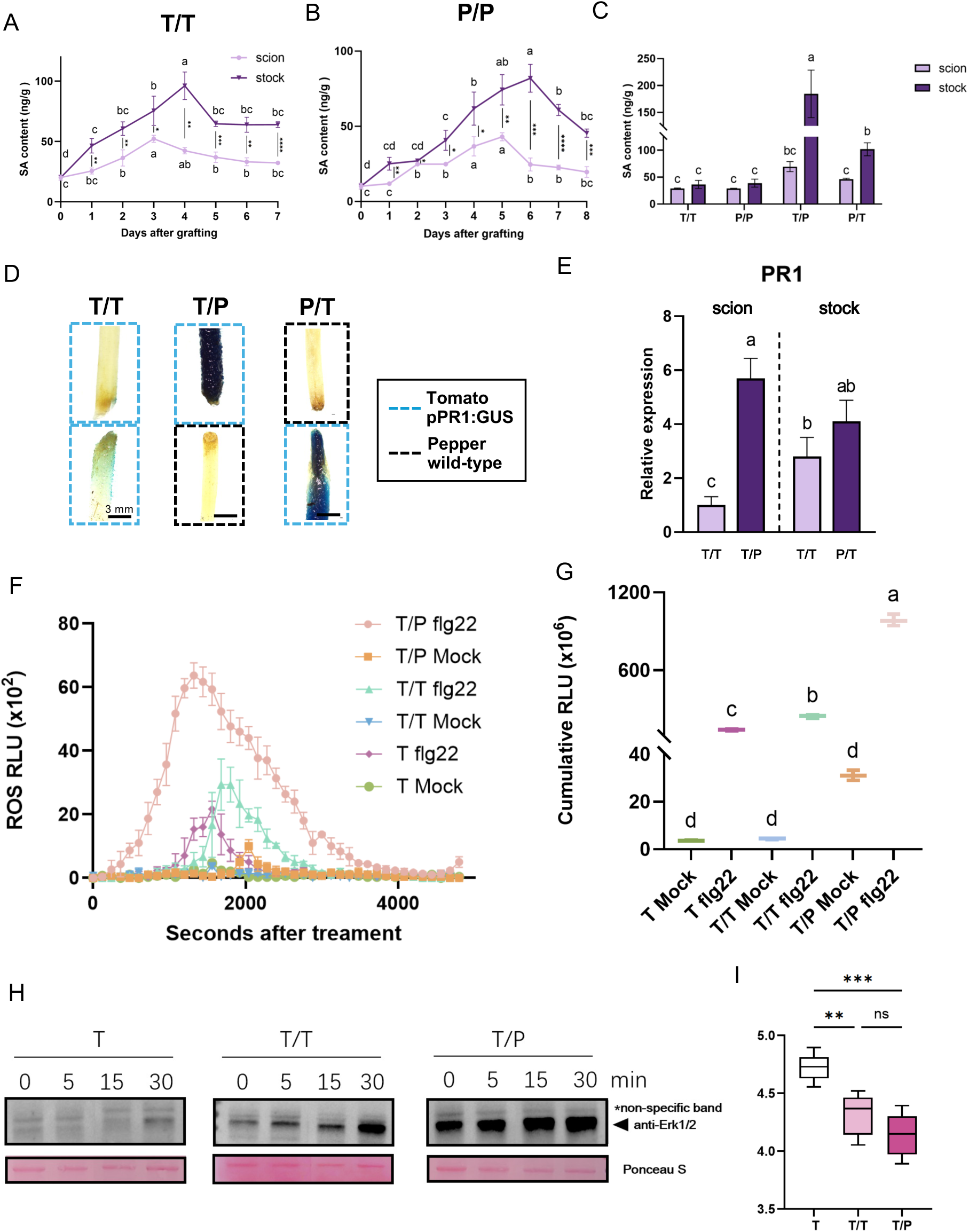
Incompatible tomato-pepper grafts overaccumulate salicylic acid during graft healing and exhibit enhanced pattern-triggered immunity (A-B) Endogenous salicylic acid (SA) concentrations in scion tissue (pink) and stock tissue (purple) during healing of tomato self-grafts (T/T; cv. Money Maker) (A) and pepper self-grafts (P/P) (B). Time is shown as days after grafting (DAG). **(C)** Endogenous SA concentrations in the scions and stocks of tomato self-grafts (T/T), pepper self-grafts (P/P), tomato/pepper heterografts (T/P), and pepper/tomato heterografts (P/T) at 7 DAG. **(D)** Representative histochemical staining of tomato cv. Alisa Craig expressing the *pPR1:GUS* reporter in self-grafts (blue box) and reciprocal heterografts with pepper (black box) at 4 DAG. Scale bars = 3mm. **(E)** Relative expression of *SlPR1* in compatible tomato self-grafts and incompatible heterografts at 7 DAG. Expression was normalized to the tomato reference gene *SlACTIN2*. Compatible grafts are shown in pink and incompatible grafts in purple. **(F)** ROS production in leaves of ungrafted tomato plants (T; cv. Money Maker), compatible tomato self-grafts (T/T), and incompatible tomato/pepper grafts (T/P) following treatment with water (mock) or 100 nM flg22. ROS production is expressed as relative luminescence units (RLU). **(G)** Cumulative RLU production over the 4800-second (80-minute) interval following mock or flg22 treatment. **(H)** MAPK phosphorylation following flg22 treatment. Leaf discs from ungrafted tomato plants (T), compatible tomato self-grafts (T/T), and incompatible tomato/pepper grafts (T/P) were treated with 100 nM flg22 and collected at the indicated time points. Phosphorylated MAPKs were detected using an anti-phospho-ERK1/2 antibody. Ponceau S staining is shown as a total-protein loading control. **(I)** Growth of *Pseudomonas syringae* pv. *tomato* DC3000 in scion leaves of ungrafted tomato plants (T), compatible tomato self-grafts (T/T), and incompatible tomato/pepper grafts (T/P). Bacterial growth is shown as log₁₀ colony-forming units (CFU) per mm². Data are presented as the mean ± SD of at least three biological replicates. In (A) and (B), scion and stock tissues were analyzed separately by one-way ANOVA followed by Tukey’s Honest Significant Difference test. Different letters indicate significant differences among time points within each tissue (*P* < 0.05). Pairwise differences between scion and stock tissues and bacterial growth were evaluated using two-tailed Student’s *t*-tests and are indicated by asterisks: ns, not significant, \**P* < 0.05, \*\**P* < 0.01, \*\*\**P* < 0.001, and \*\*\*\**P* < 0.0001. **Alt text:** Graphs and images that show incompatible tomato-pepper grafts accumulate substantially more salicylic acid at the graft junction and exhibit stronger pattern-triggered immune responses than compatible self-grafts.

To assess whether incompatible grafts exhibited differences in SA accumulation, SA content in the scion and stock was measured from compatible and incompatible grafts at 7 DAG (Figure 1C). Compatible self-grafted tomato and self-grafted pepper contained no greater than 36.4 ng/g SA, while incompatible scions (pepper/tomato, P/T; tomato/pepper, T/P) accumulated similar (ns) levels. In contrast, incompatible stocks exhibited the dominant SA response. T/P stocks showed the greatest accumulation of SA with 184.6 ng/g, a 4.8-fold increase compared to the average P/P grafts, while P/T stocks accumulated 101.9 ng/g, a 2.8-fold increase from T/T.

Focusing on the effect of SA specifically in tomato, a SA reporter line was constructed using the promoter of the SA response gene, *SlPR1* (*pPR1:GUS*; notated as *PR1*-tom henceforth). Grafted *PR1* lines were inspected 4 DAG, where peak SA accumulation was quantified in self-grafted tomato (Figure 1A), allowing for whole tissue visualization. GUS staining was detected in self-grafted tomato (*PR1*-tom/*PR1*-tom), where higher signal was detected in the stock than scion (Figure 1D). In contrast, incompatible pepper-tomato grafts (*PR1-*tom/Pepper and Pepper/*PR1*-tom) showed a dramatically increased GUS signal compared to compatible self-grafted tomato. RT-qPCR of *SlPR1* supported these data, showing that incompatible grafts elicited significantly increased *PR1* expression compared to compatible tissue 7 DAG (Figure 1E).

### Incompatible scions demonstrate increased PTI

SA is a key player in PTI and disease resistance. Since we found that incompatible grafts accumulate higher levels of SA, we wanted to test whether these grafts might mount stronger PTI responses upon pathogen perception. We first collected leaves from ungrafted (T), compatible self-grafted (T/T), and incompatible heterografted (T/P) plants 7 DAG, applied 100 nM flg22, and then profiled ROS dynamics (Figure 1F). Ungrafted and compatible grafted plants showed no ROS burst under mock treatment, while incompatible plants exhibited a late, small burst, suggesting a primed immune state in T/P grafts. Upon flg22 treatment, ungrafted plants displayed a moderate ROS burst, whereas self-grafted plants had a larger but delayed-onset response, indicating that grafting in general might modulate PTI responses in tomato.

Incompatible T/P showed an accelerated, robust, and prolonged ROS burst, highlighting hyper-activation of immune responses in incompatible combinations. Comparison of cumulative ROS response showed that, as expected, flg22 treatment elicited a significantly higher ROS response than mock in all samples (Figure 1G). However, the magnitude of the response in incompatible T/P was 4 times greater than in compatible self-grafted tissue. Next, we investigated MAPK activation and found that in response to flg22, incompatible T/P leaves showed stronger, earlier induction of MAPK phosphorylation than self-grafted leaves (Figure 1H, Supplemental Figure 1). Interestingly, compatible T/T also showed earlier, albeit less robust, MAPK phosphorylation than ungrafted tomato. This is similar to cumulative ROS, where self-grafted tomato elicited a 1.7 times greater ROS burst than ungrafted tomato (Figure 1G). Next, we conducted pathogen growth assays with *Pseudomonas syringae pv. Tomato (Pst.) DC3000* on scion leaves 7 DAG. Both incompatible T/P and compatible T/T showed increased resistance to *Pst*, compared to ungrafted plants, with T/P leaves showing the highest resistance (Figure 1I). Based on these results, we concluded that incompatible grafts have significantly upregulated PTI responses as part of the incompatibility suite, further supporting previous claims that tomato-pepper incompatibility is a side effect of overactive immunity (Thomas *et al*., 2024). Accordingly, these data suggest that, while destined for graft failure, incompatible tomato-pepper grafts can be considered immunologically primed for disease resistance. Interestingly, compatible grafted plants showed a moderate level of PTI induction, suggesting that grafting, even when compatible, induces key immune processes.

### Loss of SA accumulation cannot rescue tomato-pepper incompatible grafts

Previous work showed that incompatible tomato-pepper grafts exhibited failed vascular connections and increased cell death (Thomas *et al*., 2024). Since this work has established that incompatible grafts also exhibit high levels of SA and PTI responses, we wanted to see whether tomato plants incapable of accumulating high SA levels could improve graft compatibility when grafted onto pepper. To test this, we used the SA-deficient tomato transgenic mutant, 35S:*nahG*, which can synthesize SA but fails to accumulate it (Brading *et al*., 2000). We conducted wild-type (WT) tomato, pepper, and transgenic *nahG* tomato self-grafts, as well as reciprocal tomato-pepper and *nahG*-pepper heterografts. Self-grafted tomato (Figure 2A, H, O) and self-grafted pepper (Figure 2B, I, P) showed successful vascular reconnections at 7 DAG. As previously reported, incompatible T/P and P/T showed reduced growth (Figure 2C-D), poorly connected graft junctions (Figure 2J-K), and failed xylem reconnections (Figure 2Q-R). Trypan blue staining of the graft junctions showed that T/P and P/T also exhibited significantly more cell death in the graft junction (Figure 2X-Y) compared to compatible grafts (Figure 2V-W). Using *nahG* tomato as a graft partner with pepper did not improve graft healing, xylem connectivity, or cell death (Figure 2F-G, M-N, T-U, AA-AB), which places SA downstream of compatibility determination.

**Figure 2.**
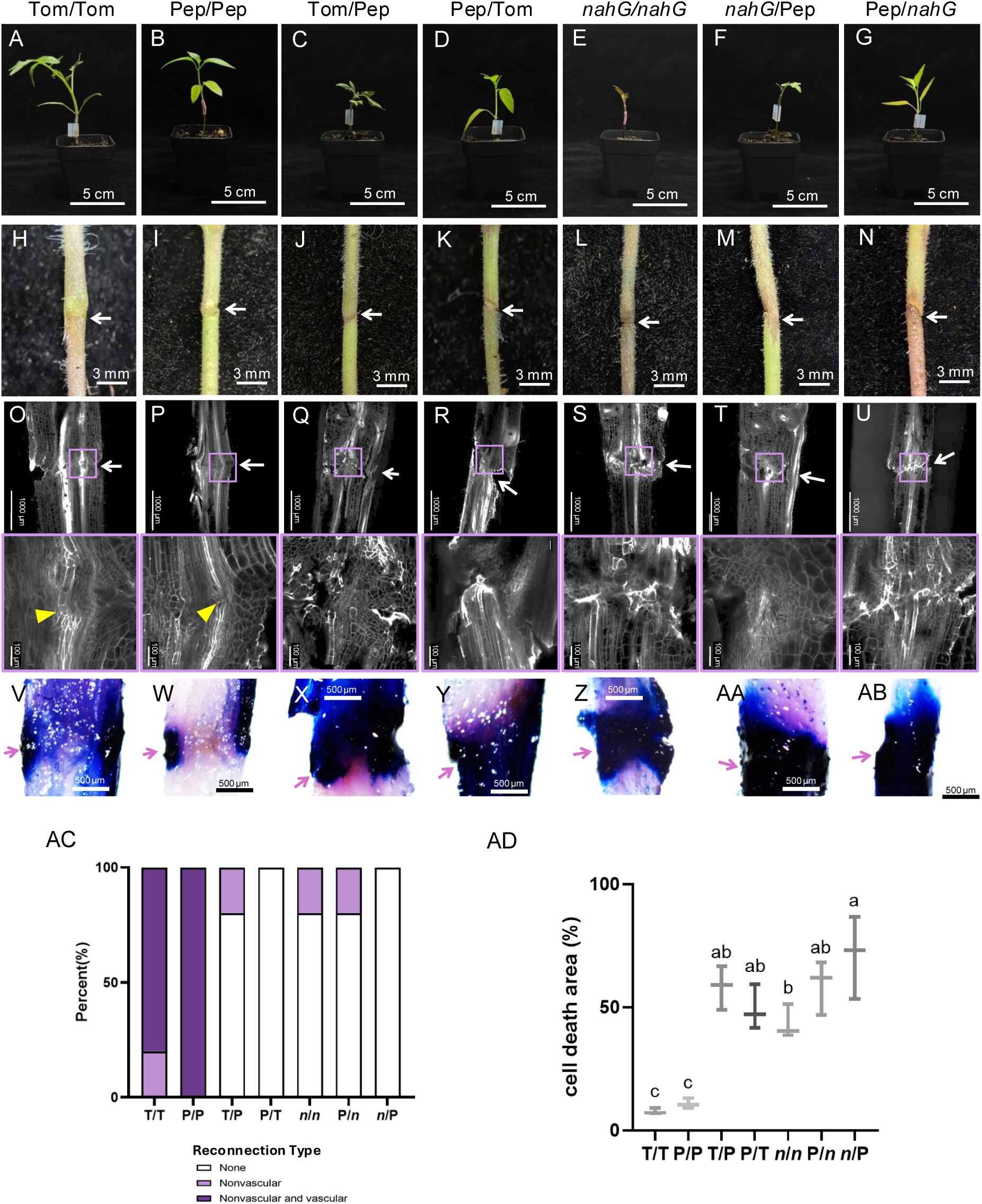
Moderate salicylic acid accumulation is required for successful graft healing (A-G) Representative images of grafted plants at 7 DAG. The graft combinations are tomato/tomato (T/T), pepper/pepper (P/P), tomato/pepper (T/P), pepper/tomato (P/T), *nahG/nahG* (n/n), *nahG*/pepper (n/P), and pepper/*nahG* (P/n). The tomato cultivar used was Money Maker. Scale bars = 5 cm. **(H-N)** Representative images of the corresponding graft junctions at 7 DAG. White arrows indicate the graft interface. Scale bars = 3 mm. **(O-U)** Representative confocal images of propidium iodide-stained graft junctions at 7 DAG. Lower panels show enlarged views of the graft interface. Yellow arrowheads indicate successful xylem reconnections. Scale bars = 1,000 µm in the upper images and 100 µm in the enlarged images. **(V-AB)** Representative light micrographs of Trypan blue-stained graft junctions at 7 DAG. Pink arrows indicate the graft interface. Scale bars = 500 µm. **(AC)** Percentage of grafts exhibiting no tissue reconnection (white), nonvascular reconnection only (light purple), or both vascular and nonvascular reconnection (dark purple). **(AD)** Proportion of the graft-junction area exhibiting Trypan blue-positive cell death as a percent of cell death area out of the total graft junction area. Quantitative data are presented as the mean ± SD of at least ten biological replicates. Differences in the proportion of cell death were evaluated by one-way ANOVA followed by Tukey’s Honest Significant Difference test. Different letters indicate significant differences (*P* < 0.05). *n* = 10. **Alt text:** Photographs, micrographs, and quantification graphs that show moderate salicylic acid accumulation is required for successful vascular regeneration and regeneration.

Interestingly, in the absence of SA (*nahG*), the self-grafted tomato healed significantly worse than WT self-grafts, with almost no fully reconnected grafts and increased cell death rates on par with incompatible T-P grafts (Figure 2E, L, S, Z). Since bench grafting is not performed under sterile conditions, grafting with this transgenic line may have predisposed the plants to graft-transmissible diseases. To test this, we allowed grafts to heal for another week and observed their phenotypes (Figure S2). However, even at 14 DAG, no obvious signs of disease were present, yet *nahG* self-grafts performed poorly, indicated by the formation of numerous adventitious roots from a scion, a common phenomenon during failed graft healing. This finding helped to sort out the order of events that occur during incompatibility, with compatibility determination occurring first, followed by increased SA accumulation. The failure of tissue to heal then likely leads to adventitious root formation and cell death at the cut site.

Since SA has previously been implicated in regulating cell expansion via XYLOGLUCAN ENDOTRANSGLUCOSYLASE/HYDROLASE enzymes (Miura *et al*., 2010; Li *et al*., 2019a; Zhang *et al*., 2025), we next profiled the expression of *SlXTHs*, which have been validated in wound and graft healing (Pitaksaringkarn *et al*., 2014; Xiong *et al*., 2026). In compatible T/T grafts, *XTH1, 2, 6, 12,* and *16* were transcriptionally abundant during healing, suggesting these genes are likely involved in the normal cell wall remodeling process of grafting (Supplemental Figure 3). In incompatible T/P and P/T, *XTHs* showed altered expression, with most genes downregulated compared to compatible grafts. In *nahG*-containing grafts (*n*/*n*, *n*/P, and P/*n*), the expression of *XTHs* was further modified, implying SA may modulate cell wall remodeling during grafting by regulating XTH enzymatic activity or expression.

To assess how the absence of SA during grafting affects the genetic program activated during compatible grafting, we performed RNA-seq on self- and heterografted tomato-pepper grafts using both WT and *nahG* tomato (Figure 3, Supplemental Figures 4-6, Supplemental Tables 1-2). We first compared all graft combinations against corresponding compatible self-grafted tissue. We found that P/*n* stock, P/T stock, and *n*/*n* stock all showed the greatest number of differentially expressed genes (DEGs; compared to T/T stock; lfc > |1.5|, adj. p-value < 0.05; Figure 3A).

**Figure 3.**
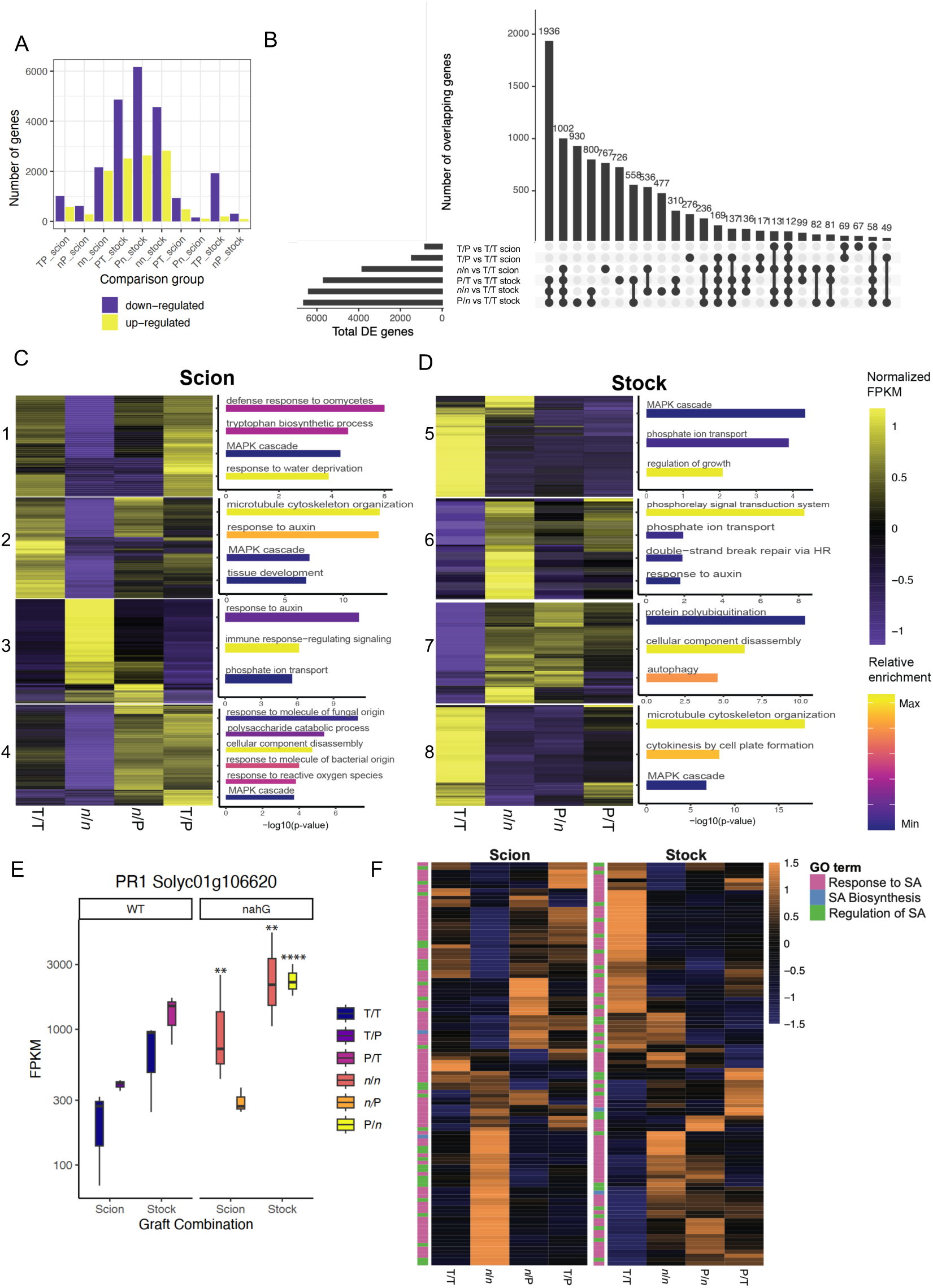
Salicylic acid acts downstream of immune activation during graft compatibility determination. **(A)** Numbers of differentially expressed genes in each graft combination relative to the corresponding tissue from compatible tomato self-grafts (T/T) at 7 DAG. Upregulated genes are shown in yellow and downregulated genes in purple. **(B)** UpSet plot showing the overlap among differentially expressed gene sets from tomato-containing graft combinations. **(C-D)** Gene-expression clusters identified by likelihood ratio testing (left) and Gene Ontology (GO) terms enriched within selected clusters (right). Each numbered module represents a distinct expression pattern. Heatmap values were scaled by row, with high expression shown in yellow and low expression in purple. In the GO enrichment plots, bar length represents statistical significance and color represents enrichment, with more highly enriched processes shown in yellow and less highly enriched processes in purple. **(E)** Expression of *SlPR1*, shown as FPKM, in tomato/tomato (T/T), tomato/pepper (T/P), pepper/tomato (P/T), *nahG/nahG* (n/n), *nahG*/pepper (n/P), and pepper/*nahG* (P/n) grafts. Differential expression was assessed relative to the corresponding scion or stock tissue of T/T grafts using an absolute log₂ fold-change cutoff of 1.5 and an adjusted *P* value < 0.05. **(F)** Row-scaled expression of genes annotated with GO terms related to SA biosynthesis, response, and regulation in T/T, n/n, n/P, T/P, P/n, and P/T grafts. High expression is shown in orange and low expression in blue. RNA-seq data were obtained from 3 biological replicates per graft combination. **Alt text:** Graphs showing transcriptomic responses for all graft combinations, supporting that salicylic acid acts downstream of immune activation during graft compatibility determination.

When comparing the overlap between all tomato DEGs, we found that a core set of DEGs was represented in all failed graft stocks (P/*n*, P/T, and *n*/*n* stocks; Figure 3B). GO term enrichment of this set of stock-specific genes highlighted metabolic reprogramming (carbohydrate metabolism and generation of precursor metabolites), defense activation (fungal response and MAPK cascade), and structural adaptations (phenylpropanoid biosynthesis and post-embryonic development) (Supplemental Table 3). In support of the unique role of SA on graft healing, we also identified a set of 726 DEGs unique to the *n/n* scion, 477 DEGs unique to the *n/n* stock, 558 DEGs only in the *nahG* self-grafts (*n/n* scion and *n/n* stock), and 131 DEGs only in *nahG* tissue (*n*/P scion, *n/n* scion, *n/n* stock, and P/*n* stock) (Supplemental Table 4-7). GO term enrichment determined that the genes DE in the *nahG* stock were upregulated for cell wall remodeling, while genes from the *nahG* scion were enriched in hormonal reprogramming via jasmonic acid and auxin.

To further explore the transcriptional patterns activated during compatible, incompatible, and SA-deficient grafting, we identified DE modules using likelihood ratio testing (LRT) and extracted GO terms of interest (Figure 3C-D, Supplemental Figure 6C-D, Supplemental Table 8-9). We found that dynamic expression patterns can be identified based on the nature of the graft and whether the tissue is scion or stock. Processes downregulated only in the *n/n* self-grafted scion included defense response to oomycetes, MAPK cascade, response to water deprivation, and tissue development (modules 1 and 2). Whereas processes downregulated in the compatible scion included responses to auxin and immune response-regulated signaling (module 3).

Furthermore, processes upregulated only in heterografted scions included responses to molecules of fungal origin and to ROS (module 4). In contrast, processes uniquely enriched in compatible stocks included MAPK cascades, regulation of growth, and microtubule cytoskeleton organization (modules 1 and 4). Processes downregulated in compatible stocks included responses to auxin and autophagy (modules 2 and 3). From this, we were able to see emerging trends where all failed grafts contained DEGs enriched for immune processes, regardless of their SA content, while compatible graft DEGs were enriched for processes involved in growth, regeneration, and auxin.

Since *nahG* appeared to show a robust immune response, we determined the expression of *PR1* in the grafted tissue (Figure 3E). In WT tissue, T/P and P/T showed increased expression of *PR1* compared to T/T scion and stock tissue (ns), while surprisingly, the *n/n* scion, *n/n* stock, and P/*n* stock all showed significantly increased *PR1* expression compared to T/T despite lacking SA accumulation. To view the global SA landscape, we extracted all genes annotated by the GO terms “response to SA”, “SA biosynthesis”, and “regulation of SA” and examined their normalized expression (Figure 3F). T/T scion showed low levels of SA processes, while T/T stock was activated. In line with the high expression levels of PR1 in *n/n*, *n/n* scions showed a *nahG*-specific SA profile, while *n/n* stocks showed trends similar to all other failed stocks. This suggests that despite lacking SA, *nahG* grafts still activate genes involved in the general defense cascade, which appear to negatively affect graft healing.

### Increased SA represses graft healing

Since a lack of SA accumulation posed novel challenges for graft healing, we next sought to determine whether exogenous SA could replicate the symptoms observed in incompatible grafts, where SA content is high. To explore this, we treated compatible self-grafted tomato with mock (water), low (0.02 mM), moderate (0.05 mM), and high levels of SA (0.1 mM), where low levels were aimed to be similar to SA levels observed endogenously, while moderate and high levels might activate processes like those in incompatible grafts. 7 DAG, all grafted plants showed comparable scion appearance (Figure 4A-D). However, high SA-treated grafts exhibited a perturbed healing response (Supplemental Figure 7A-D). We determined that exogenous SA treatment during graft healing elicited SA response genes (*NPR1* and *PRs*) and could induce endogenous SA biosynthesis (ICS and PAL pathways), as previously described (Figure 4F-J, Supplemental Figures 7E-G) (Zhang *et al*., 2010). Mock, low, and moderate SA showed typical reconnections. Whereas high SA treatment disrupted non-vascular proliferation, with 16.7% failing to adhere completely (Figure 4E, N), and triggering numerous causes (50%) of non-vascular adhesion only, proving that high levels of SA are sufficient to interfere with vascular regeneration in grafted tomato. To determine whether excess SA could also lead to increased cell death, we examined graft junctions with trypan blue at 7 DAG (Figure 4O-T). Low and moderate SA treatments did not increase cell death, but high SA significantly increased cell death at the graft junction, suggesting a gradient model for SA during grafting healing in which low-level SA promotes cell elongation and vascular reconnection, while both high and no SA blocks xylem reconnection.

**Figure 4.**
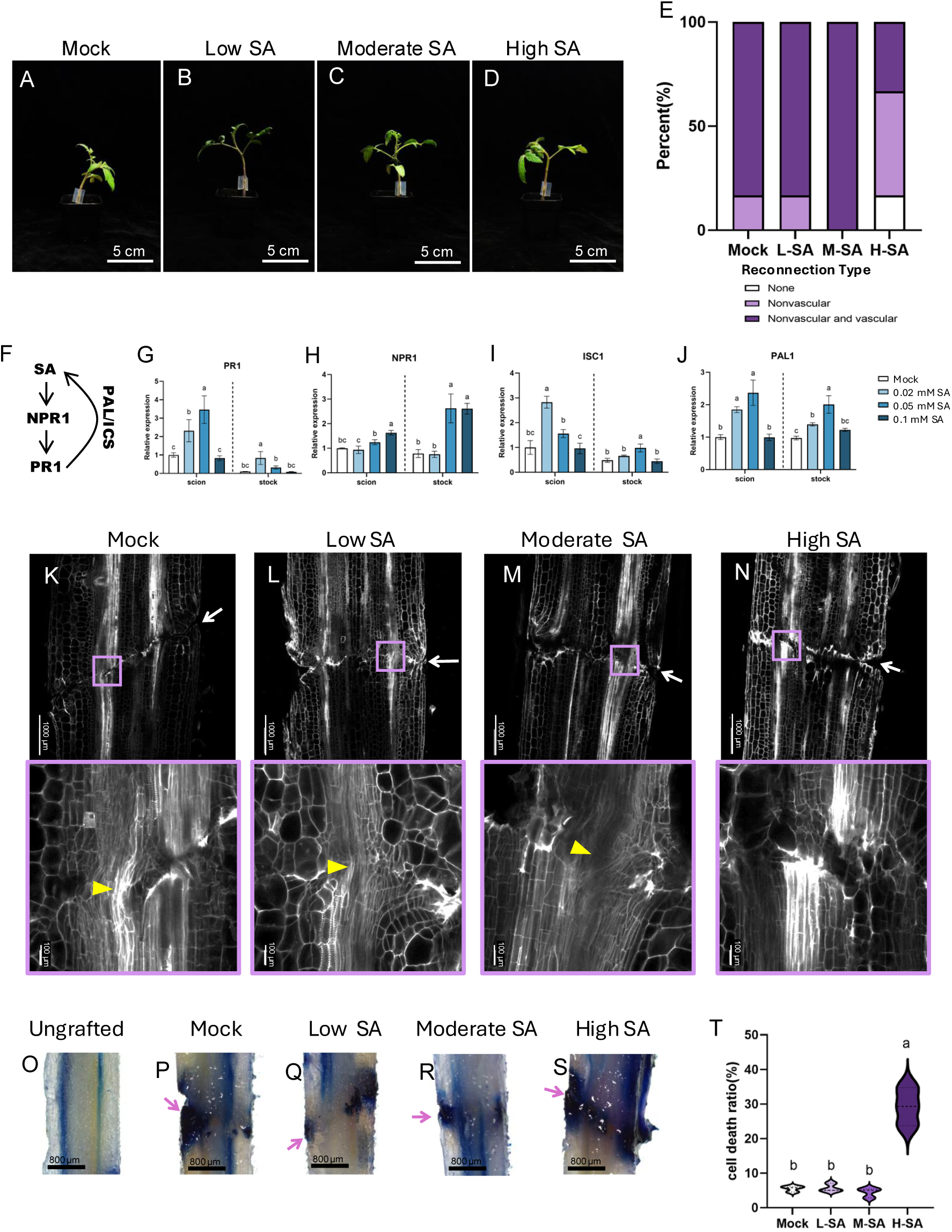
Excess salicylic acid promotes cell death and inhibits vascular reconnection in compatible grafts (A-D) Representative images of compatible tomato self-grafts treated with water (mock), 0.02 mM SA (low), 0.05 mM SA (moderate), or 0.1 mM SA (high) at 7 DAG. Scale bars = 5 cm. **(E)** Percentage of grafts exhibiting no tissue reconnection (white), nonvascular reconnection only (light purple), or both vascular and nonvascular reconnection (dark purple) following SA treatment. **(F)** Simplified model of SA biosynthesis and signaling, showing the genes analyzed in (G-J). **(G-J)** Relative expression of SA biosynthesis- and response-associated genes in compatible tomato self-grafts treated with mock, low, moderate, or high SA. Expression was normalized to *SlACTIN2*. **(K-N)** Representative confocal images of propidium iodide-stained graft junctions at 7 DAG. Lower panels show enlarged views of the graft interface. White arrows indicate the graft interface, and yellow arrowheads indicate successful xylem reconnections. Scale bars = 1,000 µm in the upper images and 100 µm in the enlarged images. **(O-S)** Representative light micrographs of Trypan blue-stained graft junctions at 7 DAG. Pink arrows indicate the graft interface. Scale bars = 800 µm. **(T)** Proportion of the graft-junction area exhibiting Trypan blue-positive cell death as a percent of cell death area out of the total graft junction area. Data are shown as violin plots. For gene-expression analyses, data represent the mean ± SD of at least three biological replicates. Scion and stock tissues were analyzed separately by one-way ANOVA followed by Tukey’s Honest Significant Difference test. For confocal and cell-death analyses, at least ten biological replicates were examined. Different letters indicate significant differences (*P* < 0.05). **Alt text**: Images, micrographs, and graphs show that excessive salicylic acid disrupts vascular reconnection and promotes cell death in otherwise compatible grafts.

### SA inhibits auxin accumulation in the graft junction

Since SA appears to be a driving factor behind the inability of incompatible grafts to establish xylem reconnections, we wanted to see if grafts containing high SA may correlate with any interesting physiological effects. To do this, we performed WGCNA on stock tissue, which showed the highest SA response, and identified the most significant cluster (ME1) (P = 9.6e-05), containing genes only upregulated in compatible grafts (Figure 5A). Within this cluster, the top GO enrichment term was response to auxin (Figure 5B). Since response to auxin was also represented in the LRT-GO analysis (Figure 3), we again extracted the genes annotated under auxin-relevant GO terms and examined their expression across all grafted tissues (Figure 5C).

**Figure 5.**
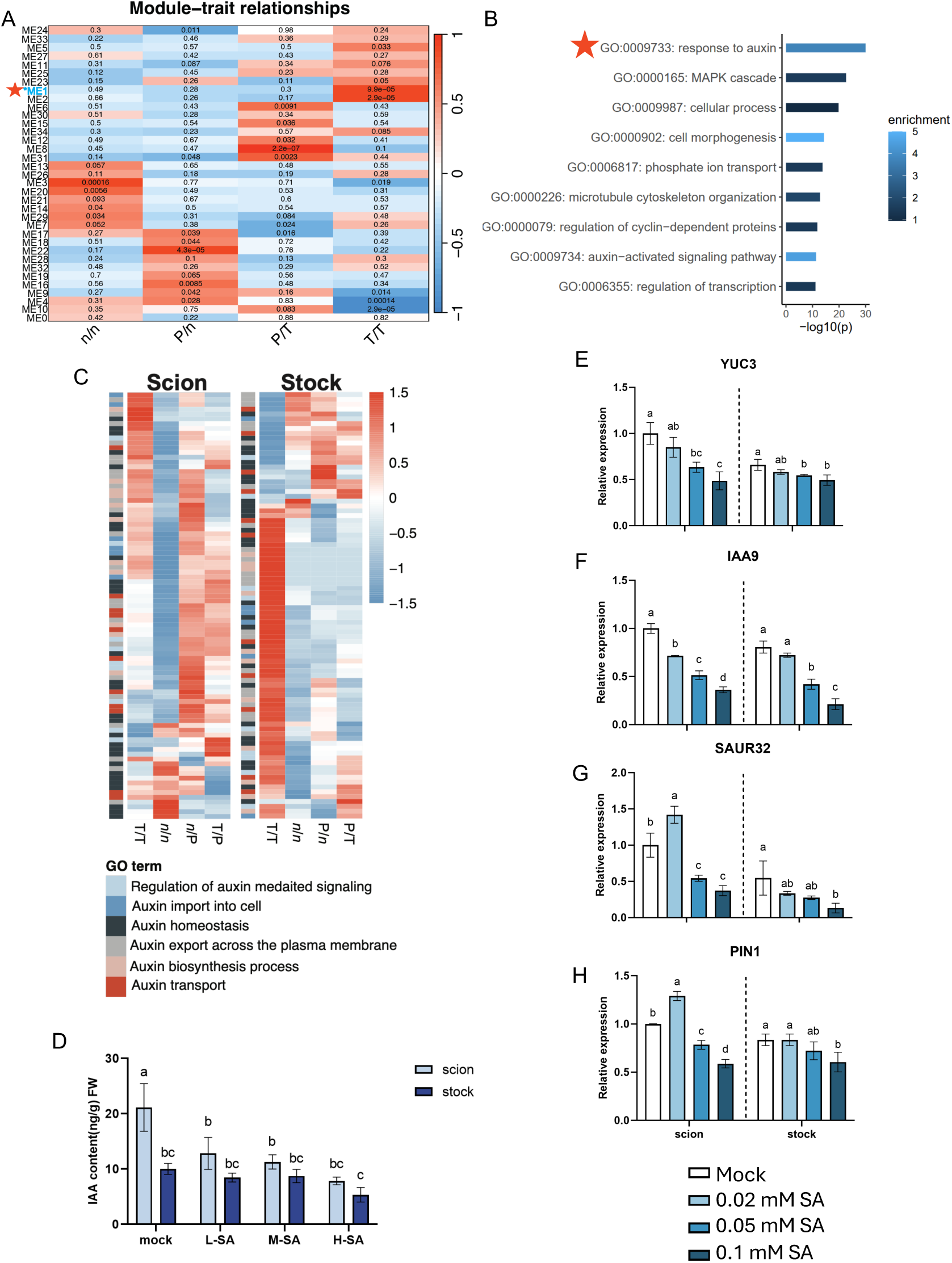
Salicylic acid suppresses auxin accumulation and auxin-responsive gene expression during graft healing. **(A)** Weighted gene co-expression network analysis (WGCNA) of tomato stock tissue from *nahG/nahG* (n/n), pepper/*nahG* (P/n), pepper/tomato (P/T), and tomato/tomato (T/T) grafts. Positive correlations are shown in red, negative correlations in blue, and correlations near zero in white. Module number (ME) is shown on the *y*-axis. Associated adjusted *P* values are displayed within each cell. The module of interest, ME1, is labeled in blue and indicated by a red star. **(B)** GO enrichment analysis of genes assigned to ME1. Bar length represents statistical significance, and color indicates enrichment, with more highly enriched terms shown in light blue. The auxin-related GO term of interest is indicated by a red asterisk. **(C)** Row-scaled expression of genes annotated with auxin-related GO terms in T/T, n/n, n/P, T/P, P/n, and P/T grafts. High expression is shown in red and low expression in blue. **(D)** Endogenous auxin concentration in scion tissue (light blue) and stock tissue (dark blue) of tomato self-grafts treated with water (mock), 0.02 mM SA (low), 0.05 mM SA (moderate), or 0.1 mM SA (high) at 7 DAG. **(E-H)** Relative expression of genes associated with auxin biosynthesis, transport, and signaling in tomato self-grafts treated with mock, low, moderate, or high SA. Expression was normalized to *SlACTIN2*. Data are presented as the mean ± SD of at least three biological replicates. For hormone and gene-expression analyses, scion and stock tissues were analyzed separately by one-way ANOVA followed by Tukey’s Honest Significant Difference test. Different letters indicate significant differences (*P* < 0.05). **Alt text:** Heatmaps and graphs that show that auxin response and accumulation are suppressed in incompatible grafts.

Compatible T/T had a completely distinct expression profile from the other graft combinations, especially in the stock, where the auxin program was practically absent from the incompatible and grafts. Interestingly, *nahG* self-grafts seemed to show a unique auxin response pattern, suggesting that perhaps *nahG* may activate an SA-independent regulation of auxin responses.

We found that exogenous SA treatment to compatible T/T (Figure 4) also led to a decrease in auxin accumulation (Figure 5D). Similarly, genes involved in auxin biosynthesis (*YUC3*), signaling (*IAA9, SAUR32*), and transport (*PIN1*) were also repressed at the graft junction with increasing SA concentration (Figure 5E-H). These findings demonstrate that SA negatively regulates auxin at the graft junction, with compatible grafts exhibiting a distinct auxin response profile that is disrupted by high SA levels or *nahG* grafts. This also helped to elucidate whether auxin responses are a consequence of failed vascular patterning or a cause, providing strong evidence for the latter.

### Incompatible grafts have reduced auxin accumulation

The effect of compatibility on auxin content was examined using the p*DR5*:GUS reporter in tomato (Figure 6A-C; DR5-tom henceforth). When compatible self-graft DR5-tom/DR5-tom was performed, GUS signal was present in the scion and stock 2 DAG, with peak expression in the scion 4 DAG, likely forming the necessary auxin maxima for vascular regeneration (Figure 6A). By 7 DAG, the scion and stock had healed, and auxin signals had returned to ungrafted levels. In contrast, DR5-tom/pepper grafts showed no GUS signal until 4 DAG, when weak auxin accumulation could be visualized (Figure 6B). Despite tissue adhesion in the DR5-tom/pepper graft, persistent GUS expression above the graft junction at 7 DAG was indicative of failed vascular reconnection and auxin pooling (Figure 6B). Pepper/DR5-tom grafts failed to elicit any GUS signal in the stock, suggesting a complete failure of PAT across the graft junction (Figure 6C). Quantification of auxin in the grafted tissue corroborated this, with T/P and P/T grafts showing significantly reduced auxin levels, specifically in the stock tissue (Figure 6D).

**Figure 6.**
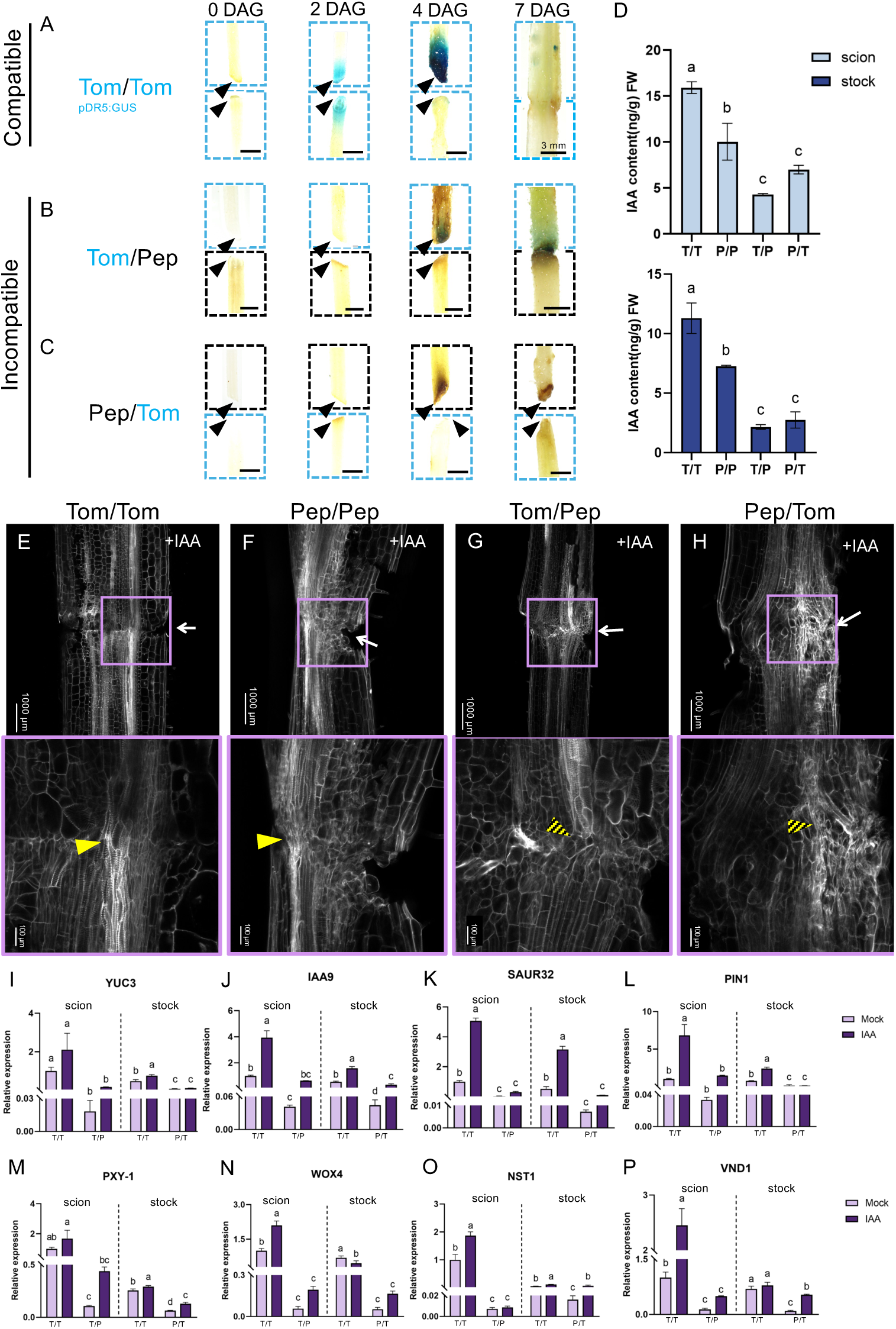
Incompatible grafts have reduced auxin accumulation, but restoration of the auxin program can partially restore graft compatibility (A-C) Representative histochemical staining of the auxin-responsive p*DR5:GUS* reporter (blue boxes) in tomato (cultivar Alisa Craig) grafted to wild-type pepper (black boxes). Reporter activity is shown in tomato self-grafts (Tom/Tom) (A), tomato reporter scions grafted onto pepper stocks (Tom/Pep) (B), and pepper scions grafted onto tomato reporter stocks (Pep/Tom) (C) at 0, 2, 4, and 7 DAG. Black arrowheads indicate the graft interface. At time points when the scion and stock remained physically separated, the tissues are shown separately; after adhesion, the complete graft junction is shown. Scale bars = 3mm. **(D)** Endogenous auxin concentrations in scion tissue (light blue) and stock tissue (dark blue) of tomato self-grafts (T/T; cv. Money Maker), pepper self-grafts (P/P), tomato/pepper heterografts (T/P), and pepper/tomato heterografts (P/T) at 4 DAG. **(E-H)** Representative confocal images of propidium iodide-stained graft junctions following exogenous auxin (100 μM IAA) treatment at 7 DAG. Corresponding mock-treated grafts are shown in Figure 3. Lower panels show enlarged views of the graft interface. White arrows indicate the graft interface, yellow arrowheads indicate complete xylem reconnection, and yellow-and-black hatched arrowheads indicate partial vascular reconnection. Scale bars = 1,000 µm in the upper images and 100 µm in the enlarged images. *n* = 10. **(I-L)** Relative expression of auxin biosynthesis-, transport-, and signaling-associated genes in compatible tomato self-grafts (T/T) and incompatible T/P and P/T grafts treated with mock solution or exogenous IAA. **(M-P)** Relative expression of vascular and cambial regulators in the same graft combinations and treatments. For (I-P), expression was normalized to *SlACTIN2*. Data are presented as the mean ± SD of at least three biological replicates. Scion and stock tissues were analyzed separately by one-way ANOVA followed by Tukey’s Honest Significant Difference test. Different letters indicate significant differences (*P* < 0.05). **Alt text:** Images, micrographs, and graphs that show that incompatible tomato-pepper grafts accumulate less auxin at the graft junction than compatible self-grafts, but that exogenous auxin can partially improve compatibility in tomato-pepper.

### Exogenous auxin has limited, but significant, effect on incompatible graft healing

Lastly, to determine if auxin alone could rescue failed xylem reconnection in tomato-pepper grafts, we applied mock (Figure 2) or supplemental auxin (100 μM) to self-grafted tomato and pepper as well as incompatible tomato-pepper heterografts (Figure 6). Unsurprisingly, compatible self-grafts showed no visible improvement in healing (Figure 6E-F), but RT-qPCR showed that auxin-responsive genes were upregulated compared to mock treatment, validating exogenous hormone perception (Figure 8I-L). In contrast, incompatible grafts appeared to show improvements in graft recovery, as evidenced by enhanced levels of nonvascular tissue and newly differentiated, but not yet cohesive, vascular tissue in the graft junction (Figure 6G-H).

RT-qPCR showed that excess auxin did not significantly activate the auxin program, with only non-significant increases observed in auxin signaling (*IAA9* and *SAUR32*), which might point to the partial rescue of downstream processes by auxin treatment (Figure 6I-L). We next determined the effect of auxin on vascular regulators and found that the addition of auxin led to significant increases in vascular regulators in compatible scions, but to only minor improvement in the stock. In contrast, incompatible grafts showed non-significant, but subtle increases in genes involved in xylem differentiation, but no improvement in cambial regulators (Figure 6M-P). This aligns with microscopic observations, which show only minor improvements to vascular healing in incompatible combinations when treated with auxin. In conclusion, while exogenous auxin treatment modestly enhanced downstream signaling, enhanced non-vascular reconnection, and partially rescued vascular development in incompatible tomato-pepper grafts, its effects were inconsistent and insufficient to fully restore functional xylem reconnection, highlighting that exogenous auxin alone cannot overcome the deleterious effect of the incompatible genetic suite in tomato-pepper grafts.

## Discussion

SA-auxin homeostasis is critical for balancing growth and defense, yet the roles of these antagonistic hormones have not been studied during graft healing. Here, we describe the SA response during compatible self-grafted tomato, self-grafted pepper, and incompatible T-P heterografts. We show that the rootstock drives SA accumulation and that incompatible grafts have significantly greater SA content than compatible grafts (Figure 1). Incompatible grafts displayed increased PTI responses, indicative of the hyper-immune state graft compatibility has been shown to trigger (Figure 1). We determined that incompatible grafts showed failed reconnection, increased cell death (Figure 2), and expression of defense-responsive genes (Figure 3). Using the *nahG* SA-deficient mutant, we explored the transcriptional and physiological effects of SA on graft healing in tomato, with a specific focus on its effects on growth and defense. Surprisingly, we found that a lack of SA could not rescue graft incompatibility and, in fact, could inhibit graft healing in *nahG* self-grafts. This led us to hypothesize that low levels of SA, like those seen in compatible grafts, may be required for graft healing, but that excess SA could inhibit the reconnection process. To test whether excessive SA has a negative effect on grafting, we applied a gradient of supplemental SA to self-grafted tomato and determined that SA could phenocopy incompatibility at high levels, blocking healing and increasing cell death rates (Figure 4). Bioinformatic analyses showed that compatible rootstocks alone were enriched for auxin responses compared to failed grafts, and that supplemental SA could inhibit genes involved in auxin biosynthesis, signaling, and transport in self-grafted tomato (Figure 5). We then validated that incompatible grafts have less auxin accumulation and that supplemental auxin could improve graft outcomes in tomato-pepper grafts (Figure 6). This work identified SA as a key hormone during graft compatibility, where low to moderate levels of SA appear to promote healing, while excess SA inhibited graft healing by interfering with auxin, a key regulator of vascular differentiation.

### Tomato graft healing requires an optimal SA response

Successful graft healing requires rapid wound closure, vascular regeneration, and the re-establishment of communication between scion and rootstock. Although auxin has been extensively studied in these processes (Zhai *et al*., 2021; Serivichyaswat *et al*., 2024), the contribution of SA has remained largely unexplored. Here, we demonstrate that SA is not simply a marker of immunity during grafting but a regulator of graft healing. Our results build on previous transcriptomic studies of tomato-pepper incompatibility, which reported enrichment of defense-related pathways and SA-responsive genes but did not determine whether SA contributed directly to incompatibility or simply reflected downstream immune activation (Thomas et al., 2024). Our findings place SA downstream of compatibility determination, likely excluding SA biosynthesis and accumulation as logical targets for compatibility expansion.

One of the most unexpected findings of this study was that despite the negative effect of excess SA on graft healing, mutants lacking SA accumulation could not improve graft outcomes. Instead, *nahG* self-grafts exhibited failed healing despite being seemingly genetically compatible and without disease (Figure 2-3). This led to the hypothesis that low-level SA is required for the typical wound response that is activated during tomato grafting.

Additionally, despite the lack of SA in *nahG* tomato, we identified numerous immunity-related genes upregulated following grafting, such as *PR1* (Figure 3E). The activation of immune processes, even PR genes, in the absence of SA has been demonstrated before (Moon *et al*., 2015; Nie *et al*., 2017), suggesting *nahG* self-grafts might undergo an SA-independent immune response that negatively affects tomato graft healing. But it is worth noting that recent studies in *Arabidopsis thaliana* overexpressing *nahG* found that the *nahG* transgene may induce a wide array of off-target effects beyond reduced SA (Wang *et al*., 2025). In this study by Wang et al., it was found that *nahG* induced 5 times the number of DE immune genes than *sid2*, an SA biosynthesis mutant, and could induce PTI-related responses in the absence of SA or infection (2025). Given this recent finding, it is critical to interpret the results of the *nahG* graft critically, and future work should instead utilize SA biosynthesis mutants to assess more endogenous effects of SA on graft healing and compatibility.

Together, these observations suggest that successful graft healing depends on maintaining an optimal range of SA. This bimodal response is consistent with previous studies showing concentration-dependent effects of SA on plant growth. Low concentrations of SA have been reported to promote cell expansion, cell division, and stress acclimation, whereas elevated SA suppresses growth and activates defense responses (Vanacker *et al*., 2001; Abreu & Munné-Bosch, 2009; Miura *et al*., 2010; Pasternak *et al*., 2019; Wang *et al*., 2021). During grafting, moderate SA accumulation may therefore promote regeneration in the junction. Once SA exceeds a set threshold, which is likely species-dependent, defense programs might supersede growth, resulting in impaired vascular regeneration and lastly cell death.

### Immune activation suppressed the necessary auxin program required for graft healing

Furthermore, we found that compatible grafts displayed strong activation of auxin-responsive genes and accumulated auxin at the graft junction, whereas incompatible grafts exhibited reduced auxin accumulation together with repression of the auxin program, especially in the stock (Figure 5). Exogenous SA reproduced this transcriptional profile and substantially reduced graft healing in compatible self-grafts, while supplemental auxin applied to incompatible grafts could partially restore expression of vascular regulation and xylem differentiation (Figure 6). Collectively, these findings support a model in which excessive SA disrupts the auxin maximum required for xylem differentiation.

Interestingly, treatment with auxin could not rescue the expression of key cambial regulators such as *WOX4* and *PXY*. Since auxin regulates multiple stages of vascular regeneration, including cambial activation and vascular patterning, our data imply that while SA can interfere with all auxin-required processes, supplemental auxin only affects cambial activity, which is not sufficient to fully rescue xylem reconnections (Suer *et al*., 2011; Solé-Gil *et al*., 2019; Canher *et al*., 2022; Serivichyaswat *et al*., 2024).

### Immunity-regeneration homeostasis underlies graft compatibility

The antagonistic relationship between SA and auxin is increasingly recognized as a central mechanism underlying the growth defense trade-off (Pasternak *et al*., 2019; Han *et al*., 2025). During pathogen infection, elevated SA suppresses auxin signaling to limit pathogen-induced growth and redirect resources toward immunity (Glickmann *et al*., 1998). Our findings indicate that this same regulatory network operates during graft healing. However, unlike pathogen infection, successful grafting requires sustained regeneration rather than prolonged defense activation. We therefore propose that graft healing depends on maintaining an appropriate balance between regeneration and immune signaling (Supplemental Figure 9). Compatible grafts transiently activate immune-wound responses but rapidly transition toward auxin-dependent regeneration, allowing vascular tissues to reconnect. In incompatible grafts, immune signaling remains elevated, SA over accumulates, and auxin-dependent developmental programs are suppressed before reconnection can be established. This framework provides a mechanistic explanation for why incompatible grafts frequently exhibit prolonged defense responses alongside defective vascular differentiation.

Importantly, this model may extend beyond tomato-pepper graft incompatibility. Environmental stress, infection, or other conditions that trigger SA accumulation could similarly compromise graft healing by suppressing auxin-dependent regeneration. Understanding how plants coordinate these competing developmental priorities may therefore have practical implications for improving graft success under various environmental conditions.

In summary, this study identifies SA as a critical regulator of graft healing and provides evidence that successful graft formation requires a finely balanced interaction between immune signaling and regeneration, via SA and auxin. We show that moderate SA accumulation accompanies compatible healing, whereas excessive SA accumulation during incompatible grafting suppresses auxin-dependent xylem reconnection. This work clarifies that, rather than acting as the primary trigger of incompatibility, SA appears to function downstream of immune activation to redirect developmental resources away from regeneration, and that incompatible cell death is likely a consequence of failed graft healing rather than the cause. These findings establish a mechanistic link between immune signaling and vascular reconnection and suggest that maintaining auxin-SA homeostasis is a key determinant of graft compatibility. More broadly, our work supports a model in which graft success is governed not simply by the activation of regeneration pathways, but by the ability of compatible plants to exit the defense response inherently activated during wounding.

## Methods

### Plant Material

Wild-type tomato (*Solanum lycopersicum*) cultivars used were Money Maker (Tomato Genetics Resource Center, University of California, Davis) and Alisa Craig (Tomato Genetics Resource Center, University of California, Davis), and pepper (*Capsicum annuum*) cultivar Xiangla No.7 (Chunyang Seed Store, Yancheng District, Luohe City).

The *pPR1:GUS* transgenic line was generated in *Solanum lycopersicum cv.* Alisa Craig, through insertion of the 2048b promoter region of Solyc01g106620 in p1300-pBI101 by Jiangsu Sanshu Biotechnology Co., Ltd. (Nantong, Jiangsu, China). The *pDR5:GUS* reporter line in *Solanum lycopersicum cv.* Alisa Craig was acquired by M. G. Ivanchenko of Oregon University (Eugene, OR, USA). The *35S*:*nahG Solanum lycopersicum cv.* Money Maker line was acquired by Professor Xiuming Li of Shandong Agricultural University (Taian, Shandong, China).

### Plant growth conditions and grafting

Seeds were surface sterilized in 70% ethanol and germinated in water with shaking at 28°C for 3 days. Seedlings were transferred to peat, vermiculite, and perlite (2:1:1) substrate (Jiaxing Chunya Horticulture Co., Ltd.; Jiaxing, Zhejiang, China) and grown in a controlled-environment growth chamber (Zhejiang Qiushi Artificial Environment, Hangzhou, China) under a 12 h photoperiod, 25 °C/20 °C (day/night) temperature, and 400 μmol m^-2^ s^-1^ photosynthetic photon flux density (PPFD). Plants were provided with Hoagland nutrient solution at the time of watering.

Seedlings were grafted at 3 weeks after sowing using the splice grafting method as previously described (Thomas et al., 2022). All graft combinations were performed as self- and reciprocal heterografted combinations. Hypocotyls were cut at approximately a 45° angle using razor blades, and attached to the rootstock using 2 mm silicone grafting clips (Taizhou Lvbang Horticultural Supplies Co., Ltd.; Taizhou). Plants were maintained in a high-humidity environment under plastic domes with shade cloth for 3 days prior to reintroduction to the light and gradual humidity acclimatization.

### Exogenous hormone treatments

Salicylic acid (SA) (Sigma-Aldrich^®^, 1609002, Shanghai) was dissolved in 100% ethanol and diluted to final concentrations of 0.02 mM, 0.05 mM and 0.10 mM. These concentrations were selected to approximate endogenous SA concentrations observed during compatible and incompatible graft healing (Jia *et al*., 2025). SA treatments were applied immediately at the time of grafting by dipping the scion into the SA solution and reapplied every 2 days by spraying 0.1ml until collection. Mock-treated plants received the solvent alone.

Indole-3-acetic acid (IAA) (Coolaber, PH1021, Beijing) was dissolved in deionized water and diluted to a final concentration of 100 μM. Auxin treatments were applied by dipping the scion into the IAA solution, and treatment was reapplied by spraying 0.1ml every 2 days until collection. Mock-treated plants received the solvent alone.

### Salicylic acid and auxin quantification

100 mg of scion or stock tissue was collected at the indicated time points, immediately frozen in liquid nitrogen, and stored at −80 °C until analysis. Frozen samples were ground with liquid nitrogen and 1 mL of 75% methanol containing stable-isotope labels: 5 ng/ml D_4_-SA (GLPBIO, GC49480, Shanghai) or 12.5 ng/ml D_5_-IAA (Merck, 591319, Germany) was added as previously described (Xi *et al*., 2021). The mixture was incubated at 4°C for 12 hours, followed by centrifugation at 12,000g for 10 min at 4 °C. The supernatant was collected and subjected to liquid chromatography coupled with tandem mass spectrometry (UPLC-MS/MS) by the Analysis Center of Agrobiology and Environment Science, Zhejiang University, using ultra-performance liquid chromatography coupled with tandem mass spectrometry (UPLC-MS/MS) using a Zorbax XDB C18 column (150 mm × 2.1 mm, 3.5 μm; Agilent, USA). The mobile phase consisted of a mixture of solvent A (0.1% formic acid in water) and solvent B (methanol) at a flow rate of 0.3 mL min^−1^ with the following gradient: 0 to 1.5 min, A:B at 60:40; followed by 6.5 min of solvent A:B at 0:100; subsequently returning to solvent A:B at 60:40 for 5 min until the end of the run.

The column temperature was kept at 40°C, and the injection volume was 20 μL. A negative electrospray ionization mode was used for detection. Phytohormone accumulation was expressed as nanograms per gram of fresh mass leaf material. Stable isotope-labeled internal standards were included during extraction for absolute quantification.

### GUS staining

β-glucuronidase (GUS) staining was performed using a commercial GUS staining kit (Coolaber^®^, Beijing, China) as previously described (Zheng *et al*., 2025). Graft junctions or stem tissue were immersed in freshly prepared staining solution and vacuum-infiltrated five times for 2 min each to facilitate reagent penetration. Samples were incubated at 37 °C for 12 h in the dark before chlorophyll was removed using washes in 70% ethanol until tissues became transparent. Stained samples were imaged using a SZ61 light microscope (Olympus, Tokyo, Japan).

### ROS burst assay

5 mm leaf disks were collected from the youngest fully expanded scion leaves of ungrafted or grafted 7 DAG. Leaf discs were floated overnight in sterile distilled water in sterile 6-well plates placed in humidity boxes under normal growth conditions.

The following day, water was replaced with assay solution containing X luminol, X horseradish peroxidase, and 100nM flg22 peptide (Absin, abs45152926, Shanghai) or water. Luminescence was recorded immediately using a 96-plate luminometer (Hangzhou KodiCheng Biotechnology Co., Ltd., Hangzhou) at 120-second intervals for 4800 seconds. Total photon counts were calculated by integrating luminescence over the assay period as relative luminescence unit (RLU). Cumulative ROS was determined by calculating the area under the curve of RLU.

### MAPK activation assay

The same protocol as the ROS burst was used to collect and prepare leaf discs. The following day, 100 nM flg22 peptide (Absin, abs45152926, Shanghai) was applied to the discs. Samples were collected at 0, 5, 15 and 30 minutes after treatment. Total protein was extracted using extraction buffer containing 50 mM Tris-HCl (pH 8.0), 150 mM NaCl, 1 mM EDTA, 0.2% Triton X-100, 0.1% β-mercaptoethanol, PhosSTOP phosphatase inhibitor (Roche, Cat. No. 4906845001, Shanghai), Pierce protease inhibitor tablets (Thermo Fisher Scientific, Cat. No. A32965, USA), and 1 mM PMSF. Samples were vortexed for 30 s and centrifuged at 13,000 × g for 20 min at 4 °C. Equal volumes of supernatant were mixed with SDS loading buffer and separated by SDS-PAGE using One-Step Colored PAGE Gel Fast Preparation Kit (SKV-0055, Share-bio, Shanghai, China), as previously described (Zhu *et al*., 2024).

Proteins were transferred to PVDF nitrocellulose membranes (Millipore, USA), and phosphorylated MAPKs were detected using anti-phospho-p44/42 MAPK antibody (Cell Signaling Technology, Cat. No. 9101S, USA; 1:1000 dilution) followed by HRP-conjugated secondary antibody (Abmart, Cat. No. M21002L, USA; 1:3000). Protein bands were visualized using Ponceau S (Coolaber, SL1281, Beijing)

### Pseudomonas syringae growth assay

7 DAG or same-age ungrafted plants were inoculated with *Pseudomonas syringae pv. tomato DC3000* by spraying 1 mL of bacterial suspensions adjusted to 10^7^ CFU mL⁻¹ with 0.02% Silwet L-77. Leaf discs were collected at 5 days post-inoculation, homogenized in sterile water, serially diluted, and plated onto 5 g/L peptone, 3 g/L yeast extract, 2% (w/v) glycerol, 1.5% (w/v) agar with 250 µL/25 mL rifampicin plates. Colony-forming units (CFU) were determined following incubation at 28 °C for 2 days and expressed per mm².

### Trypan Blue staining

Cell death at graft junctions was visualized by Trypan blue staining as previously described (Thomas *et al*., 2024). Stem segments containing approximately 2 cm above and below the graft interface were immersed in Trypan blue staining solution consisting of 10 mL lactic acid (85%), 10 mL saturated phenol, 10 mL glycerol, 10 mL distilled water and 0.4 g Trypan blue (Sigma-Aldrich, T6146, USA). Samples were gently agitated for 1 h before excess stain was removed by rinsing briefly in ethanol. Samples were cleared overnight in fresh 100% ethanol with three solution changes before transfer to 60% glycerol. Cleared graft junctions were imaged using a SZ61 light microscope (Olympus, Tokyo, Japan). Images were analyzed by capturing 1300 by 1300 μm images on 1.2x zoom. Deep tissue cell death was extracted using the green channel in Fiji (Schindelin *et al*., 2012) and the area of cell death as a ratio of the total junction area was determined.

### Propidium iodide staining

Graft junctions were processed and stained as previously described (Thomas *et al*., 2022). Briefly, tissue was fixed in ice-cold FAA solution (50% ethanol, 10% formaldehyde, 5% acetic acid) followed by vacuum infiltration for 30 min and overnight fixation at 4 °C in fresh FAA. Samples were dehydrated and rehydrated through a graded ethanol series before staining with 20 μg mL⁻¹ propidium iodide (Acros Organics; CAS 25535-16-4, Belgium) for 1 h. Samples were washed in PBS, dehydrated again, transferred through a 1:1 ethanol:methyl salicylate solution (Shanghai Yuanye Biotechnology Co., Ltd., S24041, Shanghai), and cleared in 100% methyl salicylate for two weeks. Fully cleared graft junctions were hand sectioned longitudinally and imaged on a Zeiss LSM880 Confocal Microscope using an Argon Laser 514 nm beam (Germany).

### RNA extraction and RT-qPCR

Total RNA was isolated from frozen graft junction tissue using the RNAprep Pure Plant Kit (DP419, Tiangen, China) according to the manufacturer’s instructions. Approximately 100 mg of tissue was homogenized in liquid nitrogen using a high-throughput tissue grinder (JXFSTPRP-48L, Shanghai Jingxin Industrial Development Co., Ltd., Shanghai, China) for 30 seconds, 5 times, before extraction. First-strand cDNA was synthesized using the ReverTra Ace qPCR RT Kit (Vazyme, R223, China). Quantitative PCR reactions were performed using SYBR Green Master Mix (Vazyme, Q711, China) on a LightCycler 480 II Real-Time PCR System (Roche, Switzerland) using the following cycle: 95 °C for 3 min followed by 45 cycles of 95 °C for 10 s, 57 °C for 15 s, and 72 °C for 10 s. Relative transcript abundance was calculated using the 2^−ΔΔCt method with *SlACTIN2* (Solyc11g005330) used as the internal reference gene. Primer sequences are listed in Supplementary Table 9.

### RNA sequencing and bioinformatic analyses

Graft junction tissue was harvested at 7 DAG, immediately frozen in liquid nitrogen, and submitted to Hangzhou Tiangen Technology Co., Ltd. (China) for RNA extraction, library construction, and sequencing. Total RNA was extracted (AG21101, Accurate Biology, Shanghai) from each sample, followed by comprehensive quality assessment to ensure suitability for downstream applications. RNA libraries were constructed using the KAPA Library Quantification Kit for Illumina® Platforms (KAPA Biosystems, USA) and purified using Oligo(dT) beads (AGENCOURT AMPURE XP Kit, Beckman Coulter, USA). Libraries were pooled, and high-throughput sequencing was conducted on the NovaSeq X Plus platform (Illumina, USA) using a paired-end 150bp. Raw reads were quality-filtered using fastp (Chen *et al*., 2018), mapped separately to the *Solanum lycopersicum* (ITAG4) (Hosmani et al.) and *Capsicum annuum* (CM334) (Kim *et al*., 2014) reference genomes using HISAT2 (Kim *et al*., 2015), and assembled using StringTie (Shumate *et al*., 2022). FPKM of tomato and pepper are available in Supplemental Tables 1- 2, respectively. Raw reads were analyzed in R using Tidyverse and dplyr (Wickham *et al*., 2019, 2026; R Core Team, 2021) using DEseq2 with pairwise log fold change cut-offs of > 1.5 or < −1.5 and adjusted p-values < 0.05 (Love *et al*., 2014). Likelihood ratio testing was conducted using DESeq2, with clustering and adjusted p-values < 0.05. Likelihood ratio clusters are available in Supplemental Table 8. PCA plots, normalized read counts, and DEG counts were generated in DEseq2. Bar, dot, and line plots were generated with ggplot2 (Wickham, 2011). Upset plots were made using UpSetR (Conway *et al*., 2017). Heatmaps were generated using pheatmap and TBtools (Chen *et al*., 2020; Kolde, 2025). GO term enrichment was conducted using topGO as previously described (Thomas *et al*., 2024; Alexa & Rahnenführer, 2026). Complete GO enrichment can be found in Supplemental Table 3-7, 9. WGCNA was conducted using WGCNA, dynamicTreeCut, and fastcluster (Langfelder & Horvath, 2008; Müllner, 2013; Langfelder Peter *et al*., 2016).

### Statistics

All experiments were performed using at least three independent biological replicates unless otherwise stated. Data are presented as mean ± standard deviation.

Statistical analyses were performed using GraphPad Prism version 9. Differences between two groups were evaluated using two-tailed Student’s t-tests. Comparisons among multiple treatments were analyzed by one-way or two-way ANOVA, followed by Tukey’s multiple comparison test where appropriate. Differences were considered statistically significant at p < 0.05.

## Acknowledgements

H.R.T. was supported by a Zhejiang Provincial Natural Science grant |(LZYQ25C150001).

H.R.T. and Y.Z. were both supported by the Zhejiang Provincial Key Laboratory of Horticultural

Crop Quality Improvement Fund (111 project; B17039). We thank Xiuming Li at Shandong Agricultural University for their assistance.

## Author contributions

H.R.T. and R.H. designed the study. R.H., K.M., and X.W. carried out the experimentation.

H.R.T. and R.H. analyzed the data. H.R.T. and R.H. wrote the first draft of the manuscript. X.X., H.K., and Y.Z. aided in manuscript preparation. All authors contributed critically to the final draft and approved the publication.

## Data availability

RNA reads collected for this project have been deposited on NCBI GEO, and all supplemental data have been attached.

## Supplemental Figures

**Supplemental Figure 1.**
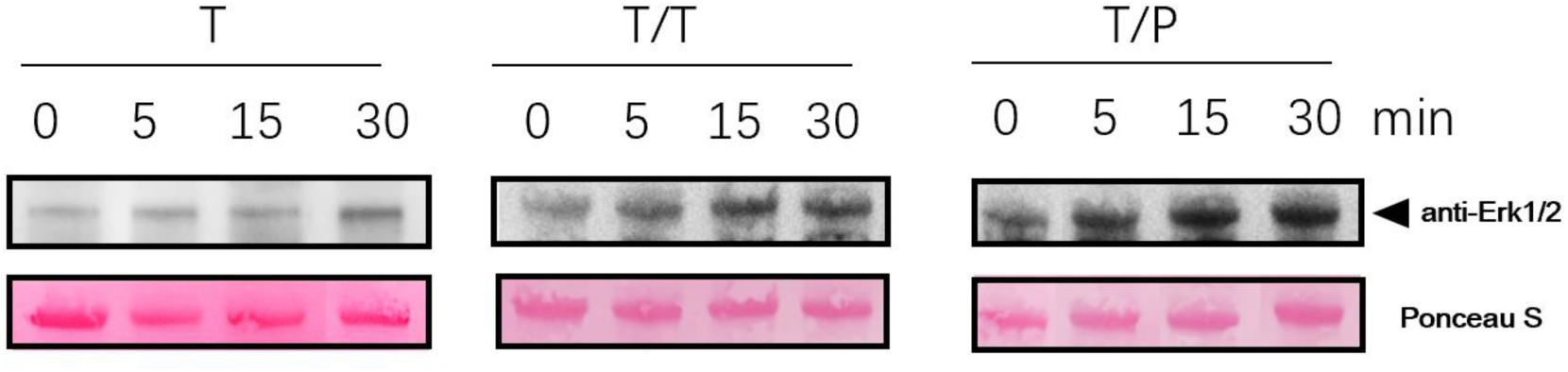
Graft incompatibility enhances flg22-induced MAPK activation. Independent biological replicate of the MAPK phosphorylation assay shown in Figure 2C. Leaf discs from ungrafted tomato plants (T), compatible tomato self-grafts (T/T), and incompatible tomato/pepper grafts (T/P) were treated with 100 nM flg22 and collected at the indicated time points. Phosphorylated MAPKs were detected using an anti-phospho-ERK1/2 antibody. Ponceau S staining is shown as a total-protein loading control.

**Supplemental Figure 2.**
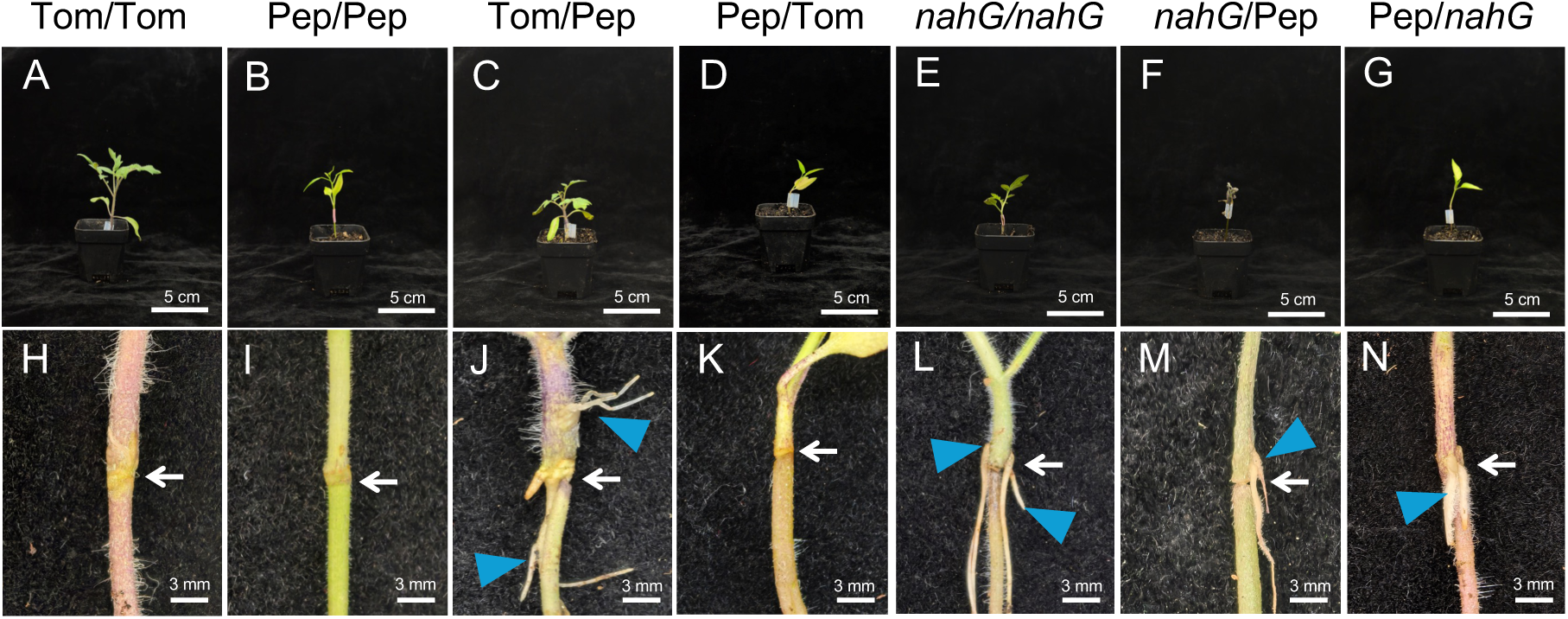
Failed grafts exhibit pronounced growth and graft-junction defects at 14 DAG (A-G) Representative images of tomato/tomato (T/T), pepper/pepper (P/P), tomato/pepper (T/P), pepper/tomato (P/T), *nahG/nahG* (n/n), *nahG*/pepper (n/P), and pepper/*nahG* (P/n) grafts at 14 DAG. Scale bars = 5 cm. *n* = 10. **(H-N)** Representative images of the corresponding graft junctions. White arrows indicate the graft interface, and blue arrowheads indicate adventitious roots arising above the graft junction. Scale bars = 3 mm.

**Supplemental Figure 3.**
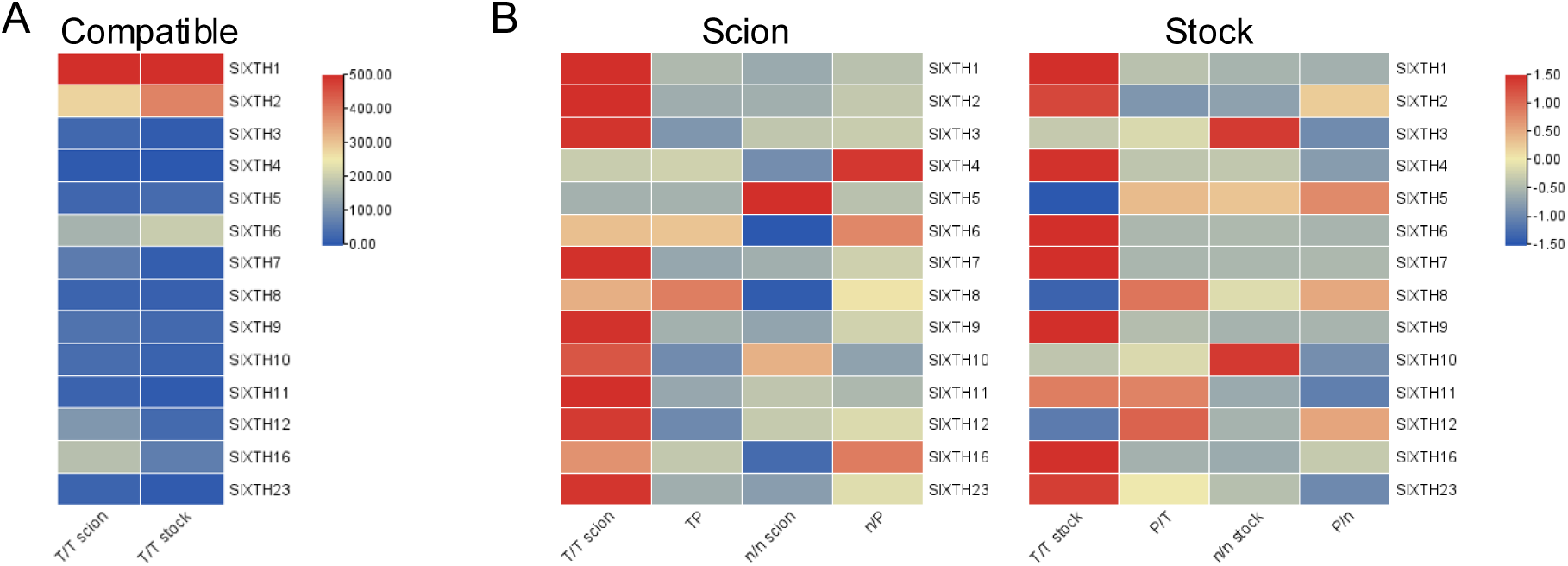
X*T*H genes are dynamically regulated during graft healing. **(A)** Expression of tomato *XTH* genes in compatible tomato self-grafts (T/T), shown as FPKM values. The displayed expression range was capped at 500, and values were not scaled. **(B)** Row-scaled expression of tomato *XTH* genes in scion tissue (left) and stock tissue (right) from the indicated graft combinations. High expression is shown in red and low expression in blue.

**Supplemental Figure 4.**
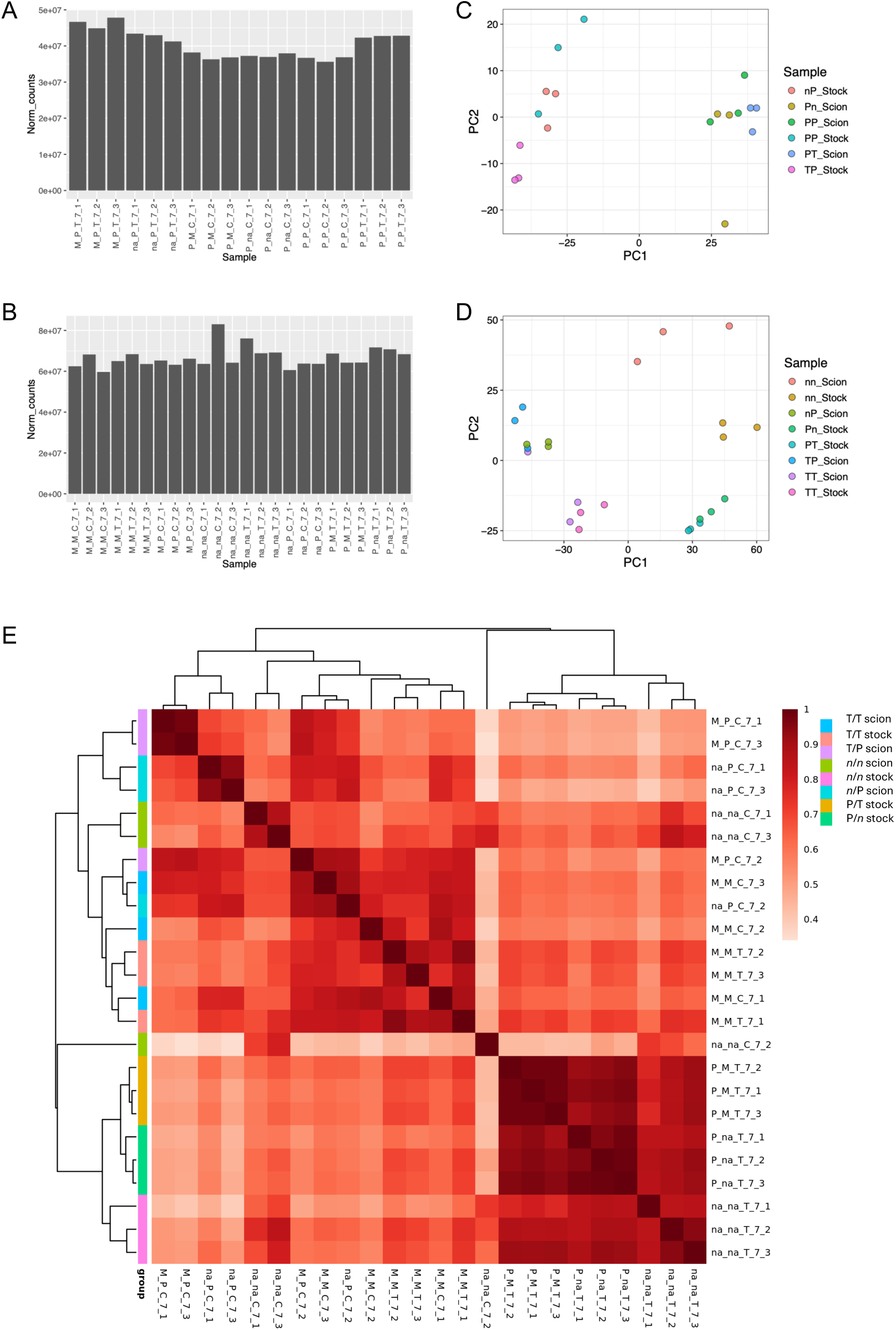
Transcriptomic profiles of *nahG* stocks resemble those of incompatible heterografts (A-B) Normalized RNA-seq read counts for tomato-derived reads (A) and pepper-derived reads (B). **(C-D)** Principal component analysis of tomato (C) and pepper (D) transcriptomes. **(E)** Sample-to-sample correlation analysis based on normalized gene expression.

**Supplemental Figure 5.**
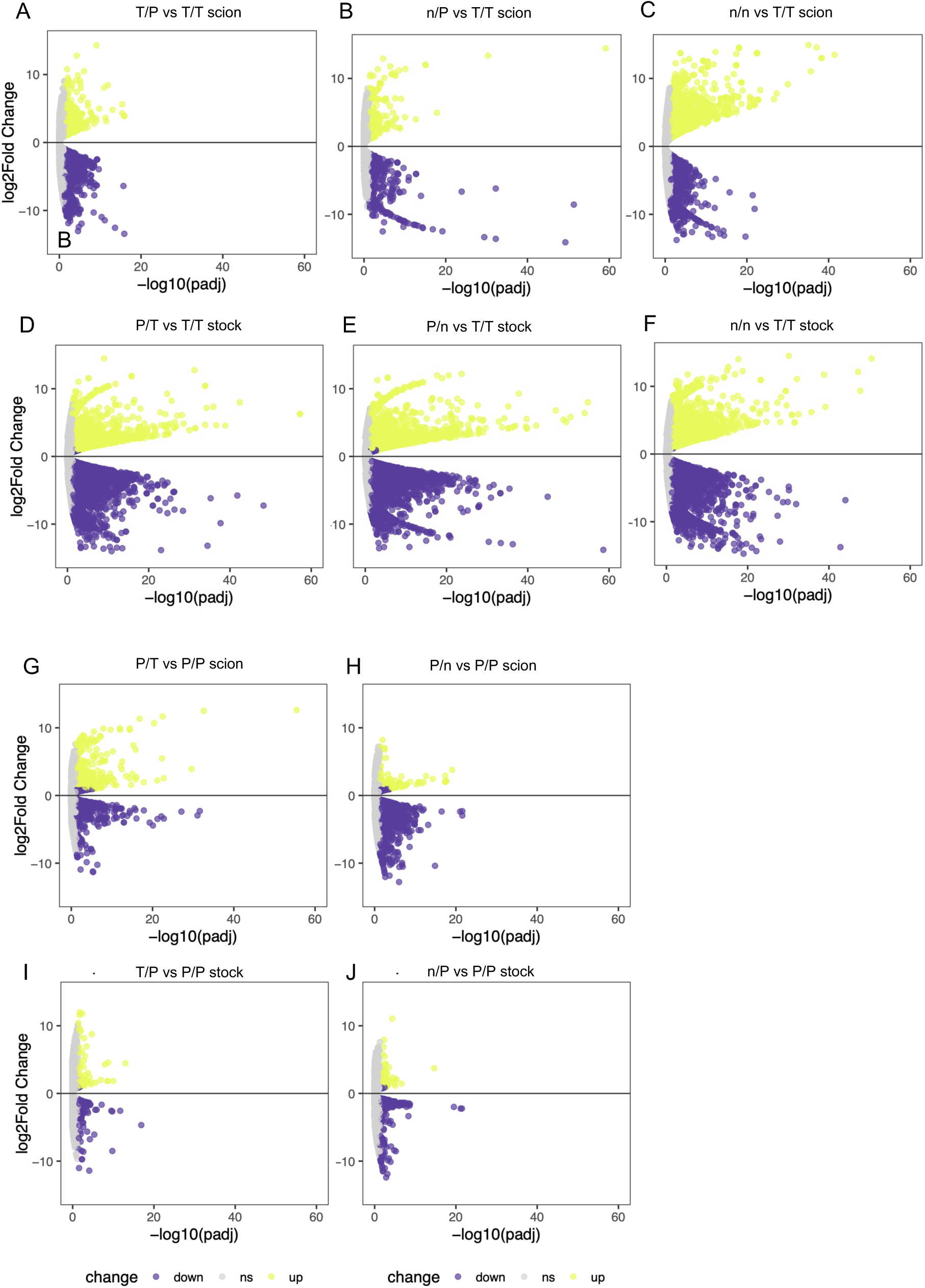
All failed grafts exhibit extensive transcriptional reprogramming relative to wild-type self-grafts (A-J) Volcano plots showing differentially expressed genes in the indicated pairwise comparisons. Upregulated genes are shown in yellow, downregulated genes in purple, and nonsignificant genes in gray. Differentially expressed genes were defined using an absolute log₂ fold-change cutoff of 1.5 and an adjusted *P* value < 0.05.

**Supplemental Figure 6.**
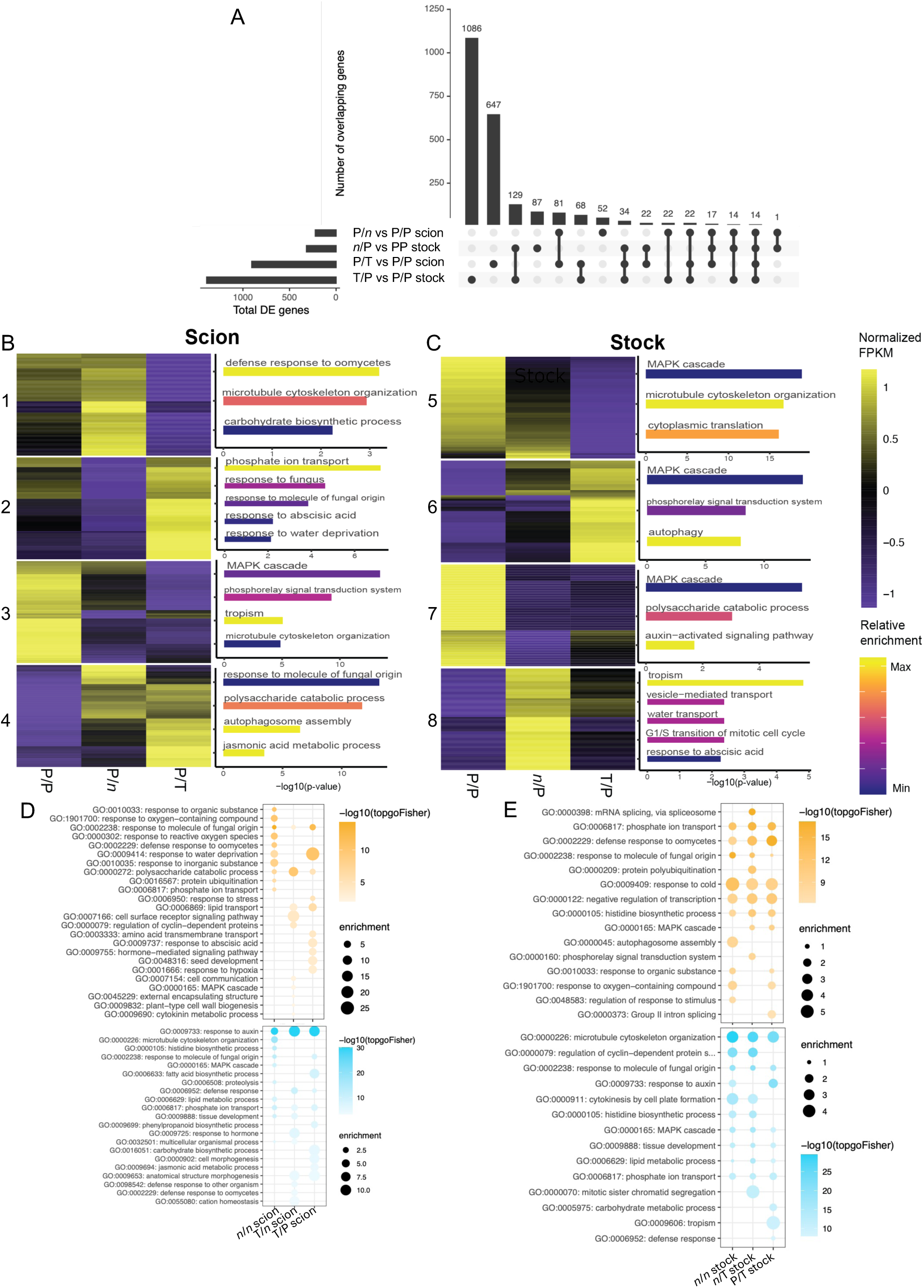
Tomato-pepper grafts exhibit reduced auxin-associated transcription relative to compatible pepper self-grafts. **(A)** UpSet plot showing overlap among differentially expressed gene sets identified in pepper tissues. **(B-C)** Pepper gene-expression clusters identified by likelihood ratio testing (left) and GO terms enriched within selected clusters (right). Heatmap values were scaled by row, with high expression shown in yellow and low expression in purple. Bar length represents statistical significance, and color indicates enrichment. **(D-E)** GO enrichment analyses of differentially expressed genes identified in pairwise comparisons with the corresponding compatible pepper self-graft tissues. Upregulated gene sets are shown in orange and downregulated gene sets in blue. Dot color represents statistical significance, and dot size represents enrichment.

**Supplemental Figure 7.**
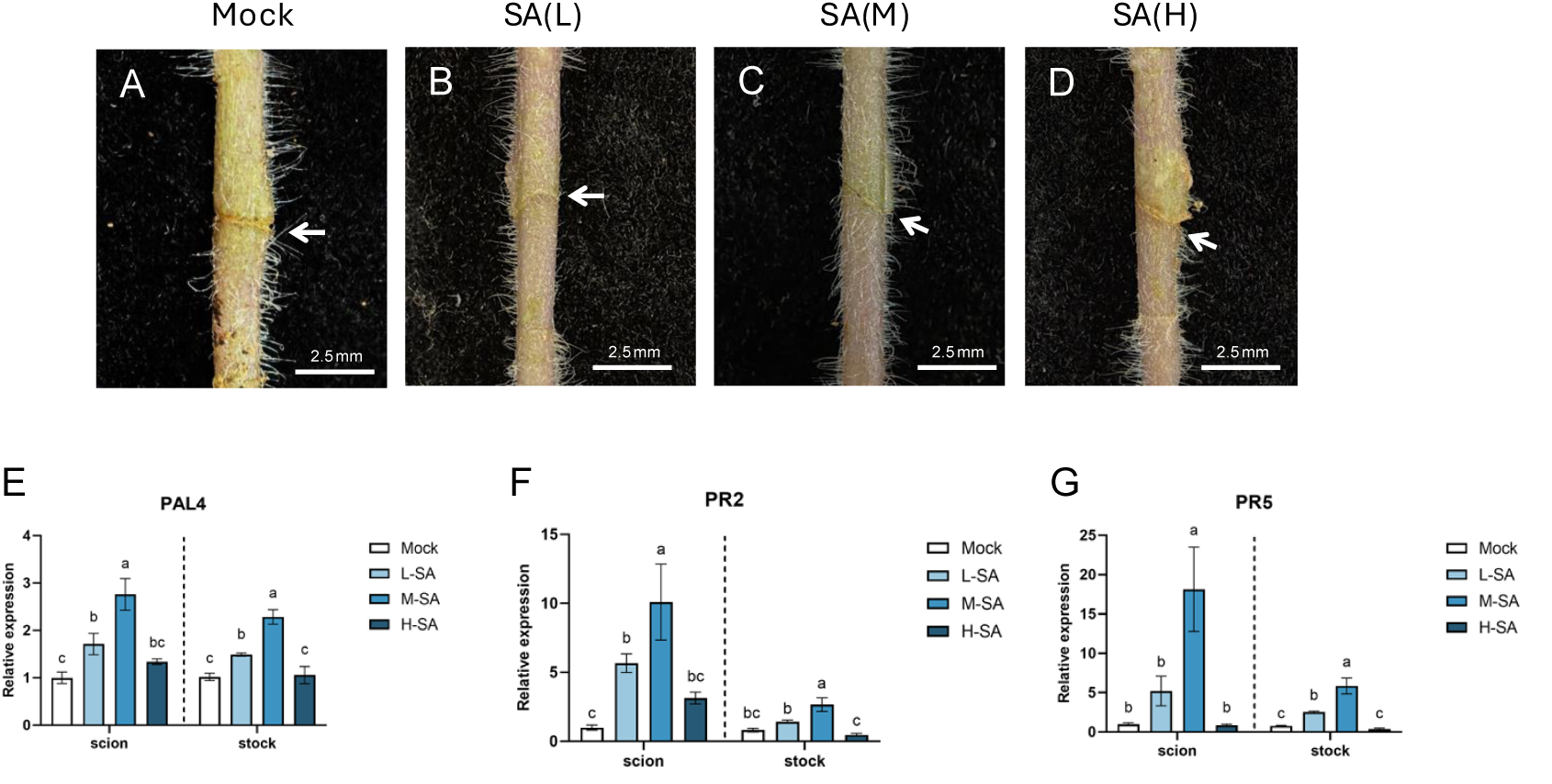
Exogenous salicylic acid activates SA-responsive gene expression at the graft junction (A-D) Representative images of tomato self-graft junctions treated with water (mock), 0.02 mM SA, 0.05 mM SA, or 0.1 mM SA at 7 DAG. White arrows indicate the graft interface. Scale bars = 2.5 mm. *n* = 10. **(E-G)** Relative expression of the SA-associated genes in tomato self-grafts following mock or SA treatment. Expression was normalized to *SlACTIN2*. Data are presented as the mean ± SD of at least three biological replicates. Scion and stock tissues were analyzed separately by one-way ANOVA followed by Tukey’s Honest Significant Difference test. Different letters indicate significant differences (*P* < 0.05).

**Supplemental Figure 8.**
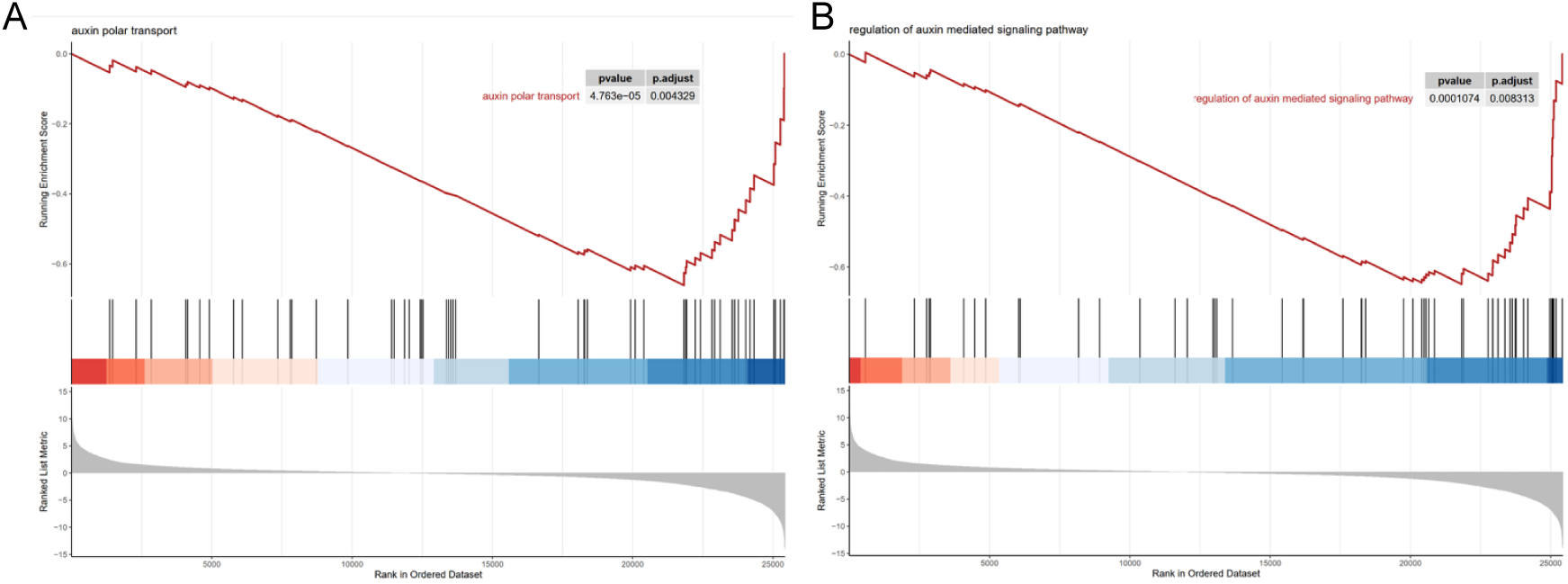
Pepper/tomato grafts exhibit reduced auxin-associated gene expression relative to compatible tomato self-grafts (A-B) Gene set enrichment analysis (GSEA) of auxin-associated pathways in pepper/tomato stocks at 7 DAG. Enrichment curves and corresponding normalized enrichment scores are shown for the indicated gene sets. Positive and negative enrichment indicate relative activation or repression, respectively, in T/T stocks vs P/T stocks.

**Supplemental Figure 9.**
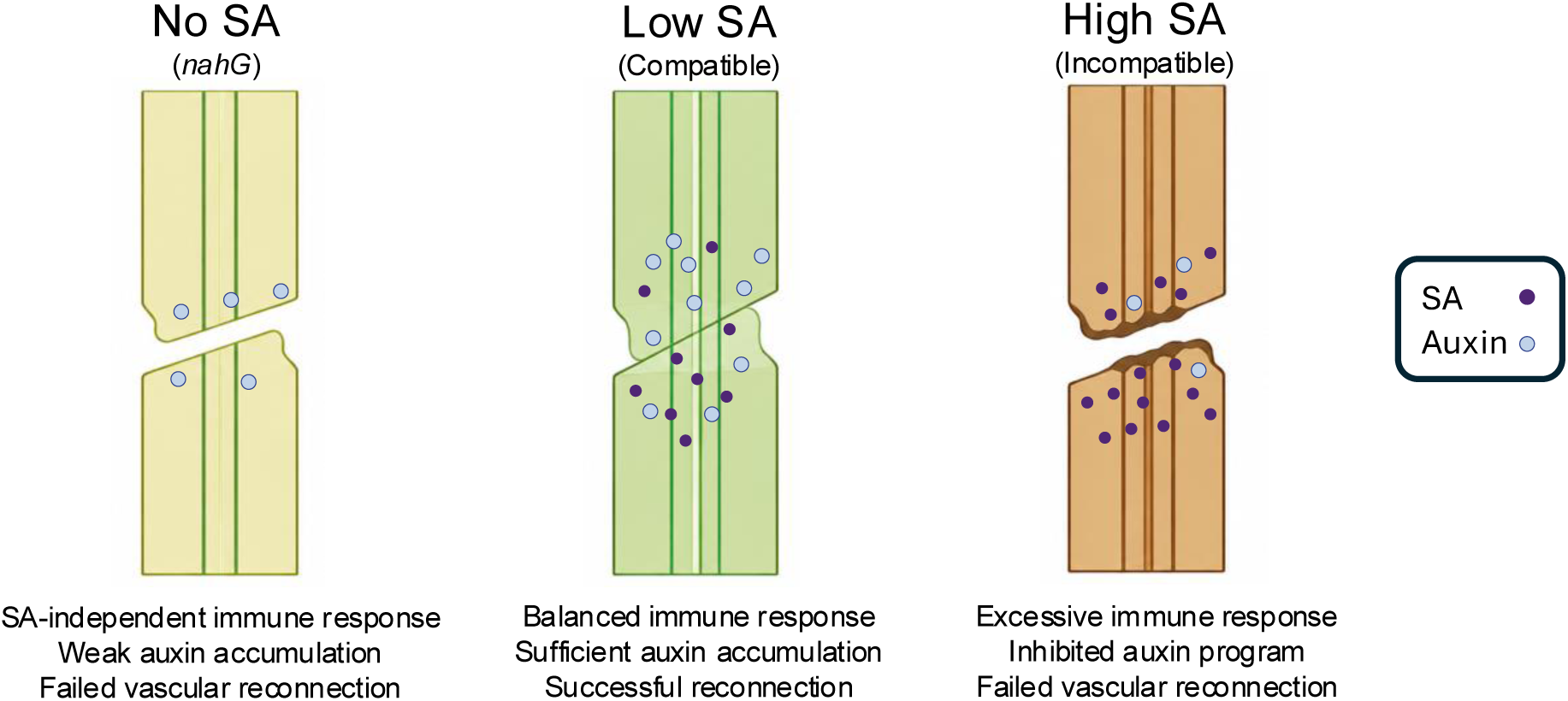
Optimal SA accumulation is required for auxin-mediated vascular reconnection in tomato grafting. Working model for the concentration-dependent role of SA during graft healing. In the absence of SA accumulation, as in *nahG* grafts, SA-independent immune responses remain active and auxin-responsive processes are reduced. Nonvascular adhesion may occur, but xylem reconnection fails. In compatible grafts, moderate SA accumulation, largely in the stock, supports basal defense while permitting auxin accumulation, nonvascular adhesion, and vascular reconnection. In incompatible grafts, excessive SA accumulation associated with sustained immune activation suppresses auxin accumulation and signaling, thereby interfering with vascular regeneration and promoting graft failure.

## Supplemental Tables

**Supplemental Table 1.** FPKM of all tomato samples 7 DAG. M = tomato, P = pepper, n = nahG, C = scion, T = stock

**Supplemental Table 2**. FPKM of all pepper samples 7 DAG. M = tomato, P = pepper, n = nahG, C = scion, T = stock

**Supplemental Table 3**. GO term enrichment for genetic overlap in genes upregulated in all failed tomato graft rootstocks (n/n, P/T, and P/nahG) compared to compatible T/T. Fisher’s adjusted p-value < 0.05.

**Supplemental Table 4.** GO term enrichment for genes upregulated in only the n/n scion compared to compatible T/T. Fisher’s adjusted p-value < 0.05.

**Supplemental Table 5**. GO term enrichment for genes upregulated in only the n/n stock compared to compatible T/T. Fisher’s adjusted p-value < 0.05.

**Supplemental Table 6**. GO term enrichment for the genetic overlap of genes upregulated in only the n/n scion and stock compared to compatible T/T. Fisher’s adjusted p-value < 0.05. **Supplemental Table 7.** GO term enrichment for the genetic overlap of genes upregulated in the n/n scion, n/n stock, n/P scion, and P/n stock compared to compatible T/T. Fisher’s adjusted p-value < 0.05.

**Supplemental Table 8**. Gene membership to likelihood ratio clusters in tomato and pepper. P-value < 0.05.

**Supplemental Table 9**. Primer sequences for RT-qPCR

